# Genome-wide analysis of DNA replication and DNA double strand breaks by TrAEL-seq

**DOI:** 10.1101/2020.08.10.243931

**Authors:** Neesha Kara, Felix Krueger, Peter Rugg-Gunn, Jonathan Houseley

**Affiliations:** Epigenetics Programme, Babraham Institute, Cambridge, United Kingdom; Babraham Bioinformatics, Babraham Institute, Cambridge, United Kingdom

## Abstract

Understanding the distribution of sites at which replication forks stall, and the ensuing fork processing events, requires genome-wide methods sensitive to both changes in replication fork structure and the formation of recombinogenic DNA ends. Here we describe <u>T</u>ransferase-<u>A</u>ctivated <u>E</u>nd <u>L</u>igation <u>seq</u>uencing (TrAEL-seq), a method that captures single stranded DNA 3’ ends genome-wide and with base pair resolution. TrAEL-seq labels DNA breaks, and profiles both stalled and processive replication forks in yeast and mammalian cells. Replication forks stalling at defined barriers and expressed genes are detected by TrAEL-seq with exceptional signal-to-noise, most likely through labelling of DNA 3’ ends exposed during fork reversal. TrAEL-seq also labels unperturbed processive replication forks to yield maps of replication fork direction similar to those obtained by Okazaki fragment sequencing, however TrAEL-seq is performed on asynchronous populations of wild-type cells without incorporation of labels, cell sorting, or biochemical purification of replication intermediates, rendering TrAEL-seq simpler and more widely applicable than existing replication fork direction profiling methods. The specificity of TrAEL-seq for DNA 3’ ends also allows accurate detection of double strand break sites after the initiation of DNA end resection, which we demonstrate by genome-wide mapping of meiotic double strand break hotspots in a *dmc1*Δ mutant. Overall, TrAEL-seq provides a flexible and robust methodology with high sensitivity and resolution for studying DNA replication and repair, which will be of significant use in determining mechanisms of genome instability.

## Introduction

DNA double strand breaks (DSBs) can be caused by exogenous agents (e.g.: ionising radiation), defective cellular processes (e.g.: replication-transcription collisions or topoisomerase dysfunction) or intentionally by the cell (e.g.: in meiosis or immunoglobulin recombination) [1–3]. We have a detailed understanding of DSB repair pathways based on decades of research [4–6], but much less understanding of which pathways are used in a given genomic context in response to particular types of damage.

Prior to the introduction of high-throughput sequencing methods, genome-wide studies of DSB formation and processing were largely restricted to meiotic recombination, where frequent DSBs at well-defined sites can be stabilised either before or after end resection and mapped on microarrays [7–9]. However these microarray methods lacked the signal-to-noise ratio required for DSB detection in other situations, and so the development of the direct DSB sequencing method BLESS marked a step-change in mapping technologies [10]. In BLESS, an adaptor is directly ligated to the DSB end to prime Illumina sequencing reads, allowing precise mapping and relative quantification of breaks. Modifications of BLESS have improved ligation efficiency (END-seq [11], DSB-capture [12]), quantitation (qDSB-seq [13], BLISS [14]), sensitivity and generality (BLISS [14], i-BLESS [15]), and variants have been developed for specific systems including meiosis (S1-seq [16]). These methods differ in detail but all involve blunting of the DNA end with nuclease activities that remove 3’ extended single-stranded DNA to form a double stranded end for adaptor ligation. This can be a problem as end resection forms extended tracts of 3’ extended single stranded DNA each side of a DSB that are degraded by blunting, such that the sequencing adaptor is ligated to the chromosomal DNA many kilobases from the original break site if resection has occurred. DNA breaks cannot therefore be mapped post-resection by BLESS-type methods, which is problematic as DSB repair is often easiest to inhibit post-resection (such as in classic *rad51*Δ or *rad52*Δ mutants in yeast).

Profiles yielded by BLESS-type methods can rarely be considered in isolation as replication has a dramatic influence on the distribution of DNA strand breaks in a cell; replication defects can be a primary cause of DNA damage but replication also provides both opportunity and the requirement to repair existing lesions. Replication forks moving rapidly through chromosomes stall at protein obstacles, DNA damage, and through collisions with the transcription machinery [17–19], and must be restarted by pathways that carry an increased risk of mutation [17–20]. Understanding the distribution and causes of DNA damage across the genome therefore requires integration of DSB profiles with approaches to monitor DNA replication.

Many methods for mapping DNA replication have been developed, which can be broadly divided into those which measure copy number changes through S-phase and those which analyse replication forks or replication bubbles directly. Copy number analysis stratifies the genome based on replication timing and defines early and late-firing origins [21–24]. This requires segregation of cell populations at different stages of replication or between replicating and non-replicating cells, either by cell-cycle synchronisation or, more flexibly, by flow cytometry. Copy number methods are well refined and the innate simplicity of this approach has even allowed application to single cells, revealing surprising uniformity in replication profiles across mammalian cells [25, 26]. However, these methods do not have the resolution to detect individual origins unless markedly different in timing, and a range of other more specialised approaches have been applied to study replication initiation [27, 28], particularly by isolating short nascent DNA strands to identify individual origins or initiation zones [29–31]. Methods have also been developed to detect replication fork directionality through isolation and sequencing of Okazaki fragments (OK-seq) [32, 33]; as well as revealing origins, these methods identify regions that are uniformly replicated in the forward or reverse direction and termination zones in which replication direction will vary depending on the point at which forks converge in individual cells. Although powerful, methods for direct analysis of forks and origins are technically demanding since replication bubbles, short nascent strands and Okazaki fragments are rare species that need to be carefully separated from each other and from contaminating genomic DNA.

Direct ligation of a sequencing adaptor to the 3’ end of individual DNA strands would be a very attractive means of quantifying DNA damage irrespective of DNA resection, and direct labelling of DNA 3’ ends may reveal replication fork direction, particularly in mutants unable to ligate Okazaki fragments. Some methods aimed at mapping single-strand breaks and base changes theoretically have this capability although they have not been applied to DSBs [34, 35], and very recently the Ulrich lab described such a method, GLOE-seq, that is capable of replication profiling in DNA ligase-deficient yeast and human cells, and also maps DSBs although activity on resected substrates was not tested [36]. Here we describe an alternative method, <u>T</u>ransferase <u>A</u>ctivated <u>E</u>nd <u>L</u>igation sequencing (TrAEL-seq), which accurately maps DNA 3’ ends at DSBs that have undergone DNA resection. Remarkably, in addition to resected DSBs we find that TrAEL-seq can profile DNA replication fork direction with excellent sensitivity even in wild-type yeast and mammalian cell populations without labelling or synchronisation.

## Results

### Overview of TrAEL-seq library construction

Addition of a short adenosine tail to DNA 3’ ends by terminal deoxynucleotidyl transferase (TdT) provides a substrate to which T4 RNA ligase can attach TrAEL adaptor 1 (Fig. 1A). This adaptor forms a hairpin that primes conversion of single stranded ligation products to double stranded DNA suitable for library construction, incorporates a biotin moiety flanked by deoxyuracil residues that allows selective purification and elution of ligation products, and includes an 8 nucleotide unique molecular identifier (UMI) for bioinformatic removal of PCR duplicates. A thermophilic polymerase with strong strand displacement and reverse transcriptase activities extends the hairpin to form un-nicked double stranded DNA, then the DNA is fragmented by sonication and adaptor-ligated material is purified on streptavidin magnetic beads. The DNA ends formed during fragmentation are polished and ligated to TrAEL adaptor 2 while still attached to the beads, then the purified fragments flanked by TrAEL adaptors 1 and 2 are eluted by cleavage of the deoxyuracil residues prior to library amplification.

**Figure 1:**
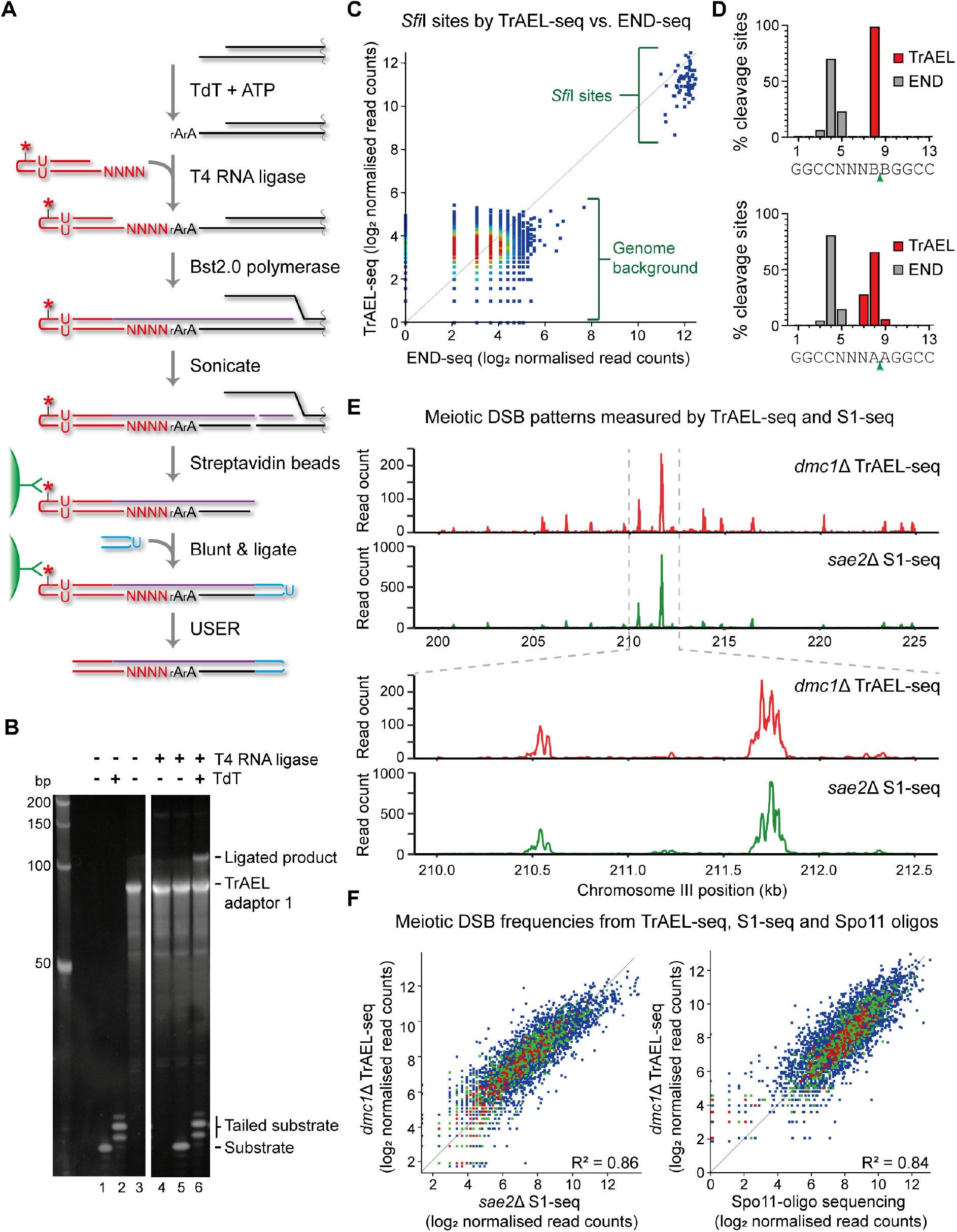
TrAEL-seq accurately maps and quantifies 3’ ends of DNA. **A**: Schematic representation of TrAEL-seq method. Agarose-embedded genomic DNA is used as a starting material, plugs are washed extensively to remove unligated TrAEL adaptor 1 and agarose is removed prior to Bst 2.0 polymerase step. The blunting and ligation of TrAEL-adaptor 2 is performed using a NEBNext Ultra II DNA kit, and TrAEL-adaptor 2 homodimers removed by washing streptavidin beads before USER enzyme treatment. The finished material is ready for PCR amplification and sequencing using the NEBNext amplification system. Note that TrAEL-seq reads map antisense to the cleaved strand, reading the complementary sequence starting from the first nucleotide before the cleavage site. * - biotin moiety, U - deoxyuracil, N - any DNA base, rA - adenosine. **B**: *in vitro* assay of adaptor ligation. An 18 bp single stranded DNA oligonucleotide was treated with or without TdT then ligated to TrAEL adaptor 1 using T4 RNA ligase 2 truncated KQ. Products were separated on a 15% PAGE gel and visualised by SYBR Gold staining. **C**: Scatter plot comparing read counts from *Sfi*I digested DNA and the genome average based on END-seq and TrAEL-seq, showing similar signal-to-noise ratio for both methods. Note that the genome average signal encompasses all single-copy 1kb regions that do not overlap with an *Sfi*I site, while *Sfi*I quantitation represents reads mapping to the 13bp recognition site. Direct comparison of read counts between the *Sfi*I and the genome average probes is misleading because of the difference in probe size. An accurate comparison of this type is not possible as the *Sfi*I cleavages are spread over 3bp (see Fig. 1D), and the majority of uncleaved 3bp sites in the genome have no mapped reads. In reality, the signal from the cleaved sites is *at least* 10 log2 units greater than an equivalent uncleaved region of the genome measured by either END-seq or TrAEL-seq. **D**: Precision mapping of *Sfi*I cleavage sites by TrAEL-seq and END-seq. *Sfi*I sites, which contain 5 degenerate bases were split into those that contain no A’s at the cleavage site (GGCCNNNB|BGGCC, 87 sites, upper panel) or A’s flanking the cleavage site (GGCCNNNA|AGGCC, 15 sites, lower panel), considering cleavage sites on forward and reverse strands separately. Mapped locations of 3’ ends were averaged across each category of site and expressed as a percentage of all 3’ ends mapped by each method to that category of site. **E**: Comparison of meiotic DSB profiles from *dmc1*Δ cells performed by TrAEL-seq and *sea2*Δ cells by S1-seq (SRA accession: SRP261135). Both techniques should map Spo11 cleavage sites in the given mutants. 25kb and 2.5kb regions of chromosome III are shown for reads counted in 20bp windows. **F:** Scatter plots of log transformed normalised read counts at all 3907 Spo11 cleavage hotspots annotated by Mohibullah and Keeney, comparing *dmc1*Δ TrAEL-seq with either *sae2*Δ S1-seq (left) or sequencing data for Spo11-associated oligonucleotides (right) [16, 42, 43, 83], (SRA accessions: SRP261135 and SRR1976210 respectively)

### Implementation of TrAEL-seq

Various ligases can, in theory, attach single stranded DNA linkers to the 3’ end of single stranded DNA but efficiency is generally poor. An alternative method recently described by Miura and colleagues utilises TdT to add 1-4 adenosine nucleotides onto single stranded DNA 3’ ends, which forms a substrate for DNA adaptor ligation by RNA ligases [37, 38]. On a test substrate *in vitro*, TdT added 1-3 nucleotide A tails to >95% of single stranded DNA molecules. This was ligated with ^~^10% efficiency to pre-adenylated TrAEL-seq adaptor 1 using truncated T4 RNA ligase 2 KQ, a ligase that is specific for 5’ adenylated adaptors (Fig. 1B).

We tested adaptor ligation using the combination of TdT and T4 RNA ligase 2 KQ at two sites in yeast genomic DNA. DNA was digested prior to ligation with the restriction enzyme *Sfi*I, then ΔC_t_ was calculated for qPCR reactions that detect TrAEL adaptor 1 ligated to an *Sfi*I cleavage site in the ribosomal DNA (rDNA) (Fig. S1A). Similarly, ΔC_t_ was calculated for TrAEL adaptor 1 ligated at the yeast rDNA replication fork barrier (RFB), a well-characterised site of replication fork stalling downstream of the 35S ribosomal RNA gene in the rDNA that can be detected by methods specific for double stranded DNA ends [13, 39] (Fig. S1A). Data from two experiments was compared with two libraries previously generated using an END-seq protocol modified for yeast (involving DNA blunting and double strand ligation, the approach of BLESS etc.). Both techniques efficiently detected these substrates, the END-seq protocol performing slightly better for the *Sfi*I break whereas TrAEL gave an equivalent or better signal for the RFB (Fig. S1B).

We then performed the TrAEL-seq procedure twice using *Sfi*I digested DNA to test the efficiency of the library synthesis steps, using qPCR to compare material sampled directly after TrAEL adaptor 1 ligation and material sampled after the whole TrAEL-seq protocol just before the final PCR amplification of the library. Remarkably, no loss of ligated material was observed across fragmentation, purification and elution steps within the detection limits of qPCR, and qPCR to detect products of the TrAEL adaptor 2 ligation gave a similar C_t_ suggesting that this step is also highly efficient (Fig. S1C) (note that this PCR product spans a heterogeneously sized population of fragments because the ligated end is the product of sonication, and qPCR is therefore only semi-quantitative). qPCR reactions performed with a single primer annealing either to TrAEL adaptor 1 or TrAEL adaptor 2 yielded no product in 40 cycles, showing that homo-adaptor dimers are not present. After library amplification, cleaning of TrAEL-seq libraries twice with AMPure beads proved sufficient to remove all heterodimers formed by TrAEL adaptor 1 and TrAEL adaptor 2 without gel purification (which is required for some prominent BLESS-type methods [11, 16]) (Fig. S1D). Library preparation through the TrAEL-seq pipeline is therefore highly efficient and shows minimal loss of material.

### Detection of 3’ extended DNA ends by TrAEL-seq

Comparing TrAEL-seq data for *Sfi*I digested genomic DNA to an END-seq library generated from equivalently digested material shows high concordance (Fig. 1C). All 70 *Sfi*I sites in the S288C sequence can be unambiguously recognised in both datasets excepting one site in the *HIS3* gene that is absent from this strain background. Fine mapping of cleavages at the *Sfi*I recognition site GGCCNNNN |NGGCC reveals differences between the methods. END-seq, in common with other BLESS-type methods, degrades the 3’ overhang and returns a consensus cleavage location 3’ of nucleotides 4-5 of the recognition site (Fig. 1D). In contrast, TrAEL-seq can map the real cleavage site (3’ of nucleotide 8) and does so for >98% events, but only for *Sfi*I sites lacking A nucleotides adjacent to the cleavage site (i.e.: GGCCNNNB |BGGCC) (Fig. 1D top). This problem stems from the A-tails added by TdT, which cannot be distinguished from genome-encoded A’s. To reconcile this issue we devised a trimming algorithm that correctly maps the *Sfi*I cleavage site to nucleotides 7-9 in >99% of reads, even when only the most challenging sites for mapping are considered (those with the structure GGCCNNNA|AGGCC) (Fig. 1D bottom). We suggest that this overall mapping accuracy of >99% within ±1 nucleotide would be sufficient for almost all applications.

A major strength of TrAEL-seq should be the ability to map DSB sites after resection, a point in the homologous recombination process that is particularly amenable to stabilisation using mutations that prevent strand invasion. We chose meiosis as an *in vivo* model system to validate this as meiotic DSB patterns have been extremely well characterised. Meiotic DSBs formed by Spo11 are processed by Sae2 amongst other factors prior to resection, after which strand invasion into a sister chromatid is mediated by Dmc1 [40, 41]. Loss of Sae2 therefore stabilises DSBs prior to resection whereas loss of Dmc1 stabilises DSBs after resection and before strand invasion. TrAEL-seq for resected DSBs in *dmc1*Δ cells 7 hours after induction of meiosis revealed a DSB pattern very similar to that observed for unresected DSBs in an *sae2*Δ mutant mapped by S1-seq (a BLESS variant specific for meiotic recombination) (Fig. 1E). Across all hotspots for Spo11 cleavage, quantitation of DSB usage frequency by TrAEL-seq correlated well with S1-seq (R^2^=0.86) (Fig. 1F, left), and to a similar extent with Spo11-associated oligonucleotide sequencing (R^2^=0.84) (Fig. 1F, right), a much more involved method that forms the gold standard for meiotic DSB mapping [42, 43]. This shows that TrAEL-seq accurately maps endogenous resected DSBs. Importantly, meiotic recombination is unusual in that mutants are known which completely stabilise DSBs, whereas stabilising breaks post-resection is often more practical in other systems.

Overall, TrAEL-seq provides an effective method for genome-wide detection and quantitation of resected DSBs.

### High resolution mapping of stalled replication forks by TrAEL-seq

Replication forks stall at various impediments during DNA replication and stalled forks may undergo reversal or cleavage as the cell attempts to restart replication (Fig. 2A). The RFB in the rDNA of budding yeast is a classic system for studies of replication fork stalling, and is defined by the binding of the Fob1 protein in each rDNA repeat [44]. RFBs are positioned just downstream of 35S rRNA genes and prevent the passage of replication forks moving against the direction of 35S transcription that would otherwise encounter the RNA polymerase I machinery head-on [45, 46]. DSBs have been reported at the RFB based both on southern blotting and qDSB-seq (a BLESS-type method for mapping double stranded DNA ends) [13, 47, 48], and the site has long been known as a hotspot for recombination [49, 50]. To detect replication forks stalled at the RFB and test the requirement for homologous recombination in resolution of these species we prepared TrAEL-seq libraries from unsynchronised wild-type, *fob1*Δ and *rad52*Δ cells growing at mid-log phase: *fob1*Δ cells lack RFB activity whilst *rad52*Δ mutants cannot initiate strand invasion and will accumulate DSBs that are normally repaired by homologous recombination.

**Figure 2:**
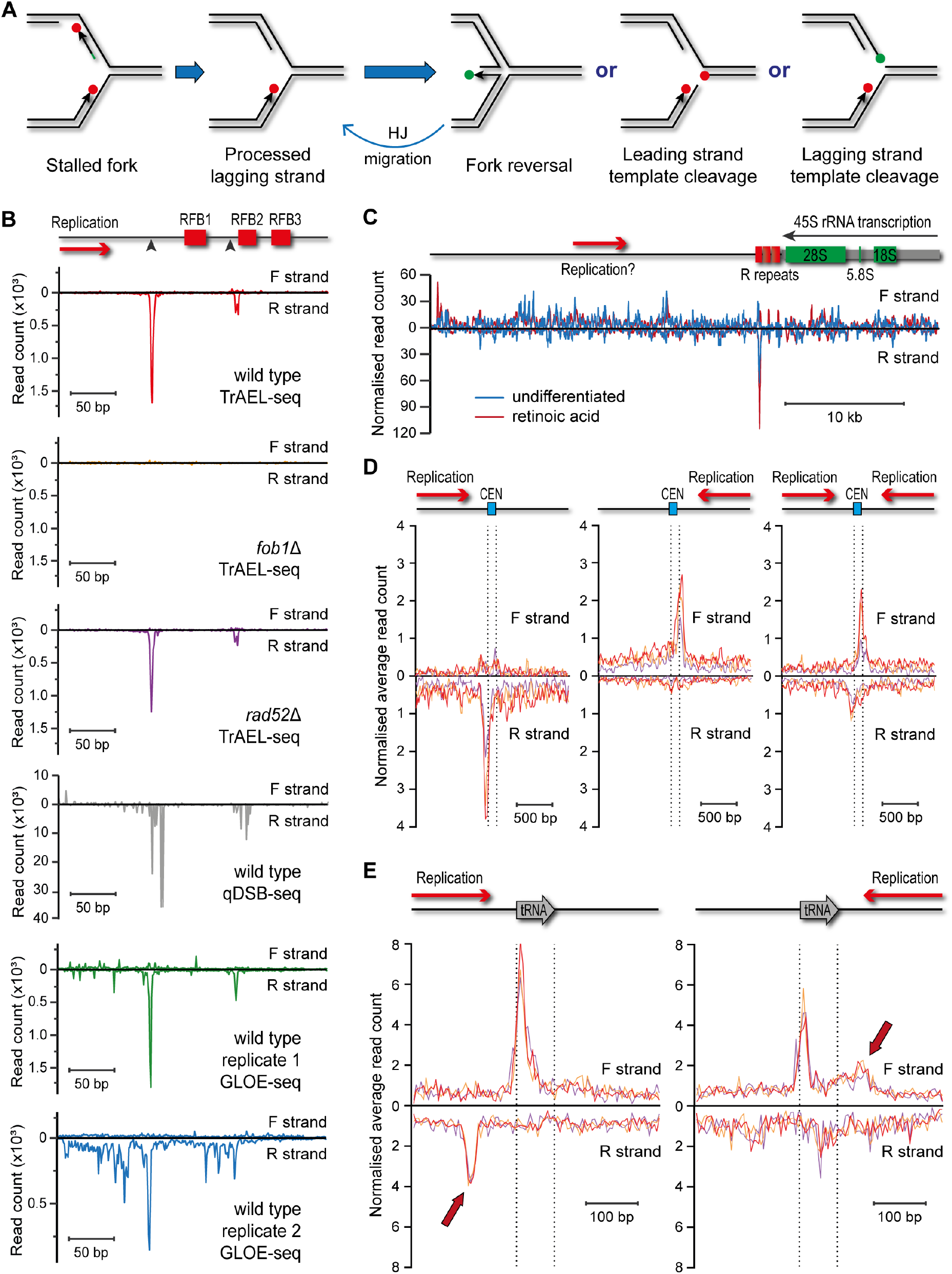
Visualisation of replication fork stalling sites by TrAEL-seq. **A**: Potential processing pathways of a stalled replication fork. Lagging strand processing is likely to finish soon after stalling, and at least for the yeast RFB it is known that the lagging strand RNA primer is removed [51]. The fork could then undergo fork reversal to yield a Holliday junction or be cleaved on the leading or lagging strand. Whereas cleavage is irreversible and requires a recombination event to re-start the replication fork, reversed forks can revert to the normal replication fork structure by Holliday Junction migration (labelled HJ migration). The various 3’ DNA ends are marked with dots, either green for those predicted to be TrAEL-seq substrates or red for those that are unlikely to be tailed by TdT. The RNA primer on the Okazaki fragment in the leftmost structure is shown in green. **B**: Comparison of the yeast rDNA RFB signals in TrAEL-seq datasets compared to qDSB-seq (SRA accession: SRX5576747) [13] and GLOE-seq (SRA accessions: SRX6436839 and SRX6436840) [36]. Reads were quantified in 1 nucleotide steps and are shown as raw read counts. Note that the three TrAEL-seq datasets are shown on the same scale and that the total number of de-duplicated reads in these libraries is very similar (range 1.68-1.97×10^6^ reads), so the absence of signal in *fob1*Δ is due to the absence of replication fork stalling rather than lower library complexity. Scales were altered for qDSB-seq and GLOE-seq datasets because of varying sequence depth and signal-to-noise ratio. qDSB-seq data was obtained from S-phase synchronised cells, all other samples are from asynchronous log-phase cell populations growing in YPD media. Schematic diagram shows the positions of RFB elements previously mapped by 2D gel electrophoresis [45, 46], and black triangles indicate previously mapped sites of DNA ends [47, 51]. **C**: rDNA TrAEL-seq reads for undifferentiated and retinoic acid-treated hESCs, normalised so that both samples have equivalent total reads after deduplication. Reads were summed in 100 bp sliding windows spaced every 10 bp. One rDNA repeat is shown, the RNA polymerase I-transcribed 45S RNA is shown as a grey line with mature rRNAs marked in green in the schematic diagram. Note that the 45S gene is shown as transcribed right-to-left to maintain consistency with the yeast data. The R repeats, which contain the RFBs, are marked in green, while the primary direction of replication is shown by a red arrow labelled as ‘Replication?’ to take into account evidence that forks can move in both directions through the human rDNA. **D**: Average TrAEL-seq profiles across centromeres +/- 1kb for wild type (red), *fob1*Δ (orange) and *rad52*Δ (purple) cells. Centromeres are categorised based on replication direction in the yeast genome assembly into those replicated forward (CEN3, CEN5, CEN13), reverse (CEN11, CEN15, CEN10, CEN8, CEN12, CEN9) and those in termination zones that could be replicated in either direction (CEN14, CEN16, CEN1, CEN4, CEN7, CEN6, CEN2), see Fig. S2A for details. **E:** Average TrAEL-seq profiles across tRNAs +/- 200 bp for wild type, *fob1*Δ and *rad52*Δ cells. tRNAs are categorised into those for which transcription is co-directional with the replication fork and those for which transcription is head-on to the direction of the replication fork. tRNAs for which the replication direction is not well defined were excluded. Arrows indicate peaks that are dependent on replication direction.

Two RFB sites are readily visible in wild type TrAEL-seq data as peaks of antisense reads relative to the direction of replication, but are absent in the *fob1*Δ mutant (Fig. 2B, wild type and *fob1*Δ panels). Both sites correspond well with previous mapping by high resolution gels [47, 51] and are also visible in published qDSB-seq and GLOE-seq datasets, although TrAEL-seq data displays higher signal-to-noise ratios than GLOE-seq data with peaks that correspond more closely to known sites than qDSB-seq peaks (Fig. 2B) [13, 36]. It should be noted that the TrAEL-seq and the GLOE-seq datasets used for this analysis derived from asynchronous cells whereas the cells for qDSB-seq had been tightly synchronised in S phase, underlining the high sensitivity of TrAEL-seq for stalled replication forks.

The structures resulting from stalled fork processing have multiple 3’ ends that could be labelled by TrAEL-seq (Fig. 2A, red and green dots), however TdT will strongly favour reversed forks and forks cleaved on the lagging strand that possess 3’ extended single-stranded ends [52] (Fig. 2A, green dots). In both cases this 3’ end is part of a double stranded end that would also be a substrate for qDSB-seq, explaining why both methods detect RFB peaks. Importantly, no difference was observed between the *rad52*Δ signal and the wild type, showing that these double stranded ends are not normally processed by the homologous recombination machinery (Fig. 2B compare wild type and *rad52*Δ panels). DSBs formed in the rDNA are known to be repaired by homologous recombination, and although we and others have reported Rad52-independent recombination at the rDNA these are rare events unknown in wild type cells [53–55]. Therefore, if these TrAEL-seq and qDSB-seq peaks represented fork cleavage events we would expect a strong stabilisation in the *rad52*Δ mutant that we do not observe, so we consider that the vast majority of the double stranded DNA ends at the RFB represent reversed forks that can revert to normal replication fork structures by Holliday Junction migration without recombination (see Discussion).

To determine the applicability of TrAEL-seq to mammalian cells, we generated TrAEL-seq libraries from 0.5 million human embryonic stem cells (hESC) that were either undifferentiated or subjected to retinoic acid-induced differentiation. A single peak was observed in the rDNA downstream of the RNA polymerase I termination site in both hESC samples, only on the antisense strand and located at the most distal of the known RFB sites (Fig. 2C) [56]. This observation is consistent with an efficient polar RFB located just downstream of the RNA polymerase I transcription unit, as seen in diverse species from plants to yeast to mice [45, 57–60]. In contrast we see no evidence of the low efficiency bi-directional RFB that has been reported in human cells (Fig 2C) [56, 61, 62]; it may be that the signal-to-noise is too low to detect fork stalling on the sense strand, but alternatively this may reflect the fact that our study was performed on unmodified hESCs, whereas the previous human studies have been performed on transformed cell lines.

rDNA RFBs are not the only sites at which replication forks stall, for example reported GLOE-seq peaks at yeast centromeres likely stem from forks stalling at centromeric chromatin [36, 63]. To probe this relationship we first stratified centromeres into those replicated only by reverse forks, those replicated only by forward forks and those sited in termination zones where forks converge (Fig. S2A). At centromeres replicated from one direction only, we observed an accumulation of reads antisense to the direction of replication just before the centromere, while forks in termination zones that can be replicated in either direction displayed both peaks (Fig. 2D). A similar analysis of tRNA loci, which are also known to stall replication forks [64], yielded more complex patterns (Fig. 2E). These regions displayed antisense peaks upstream or downstream of the tRNA depending on the direction of replication, consistent with previous studies that reported both co-directional and head-on tRNA transcription can stall replication forks, at least in the absence of replicative helicases (Fig. 2E arrows) [65–67]. However we also observed a major peak covering the first ^~^15bp of the tRNA gene, which was not affected by replication direction and appears to mark a transcription-associated break on the template strand that must be a conserved feature of tRNA transcription as it is also detected in the hESC samples (Fig. S2B). This aside, we find that sites of replication fork stalling both at the RFB and other sites are revealed by an accumulation of TrAEL-seq reads antisense to the direction of replication.

Taken together, these results show that TrAEL-seq allows highly sensitive and precise mapping of replication fork stalling, most likely through labelling of reversed replication forks.

### TrAEL-seq profiles describe replication fork directionality

A striking feature of yeast TrAEL-seq data is the massive variation in strand bias of reads in different regions of the genome: a violin plot of the fraction of reverse reads in 1 kb bins shows two distinct peaks at 20% and 80%, a behaviour only weakly observed in comparable GLOE-seq data (Fig. 3A) [36]. Read polarity forms clear domains when represented on Replication Fork Direction (RFD) plots, which show the difference between forward and reverse read counts as a function of genomic location (Fig. 3B). TrAEL-seq read polarity in asynchronous wild-type cells almost perfectly mirrors the GLOE-seq map of Okazaki fragment ends from a DNA ligase depletion experiment (*cdc9*) [36], strongly suggesting that TrAEL-seq detects processive replication forks (Fig. 3B and S3A). Notably, the locations at which TrAEL-seq polarity switches from negative to positive coincide with known replication origins (Autonomously Replicating Sequences or ARS) (Figure 3B dotted vertical lines), and alignment of TrAEL-seq reads across 30 kb either side of all ARS elements reveals a precise switch in polarity as would be expected for replication forks diverging from replication origins (Figure 3C).

**Figure 3:**
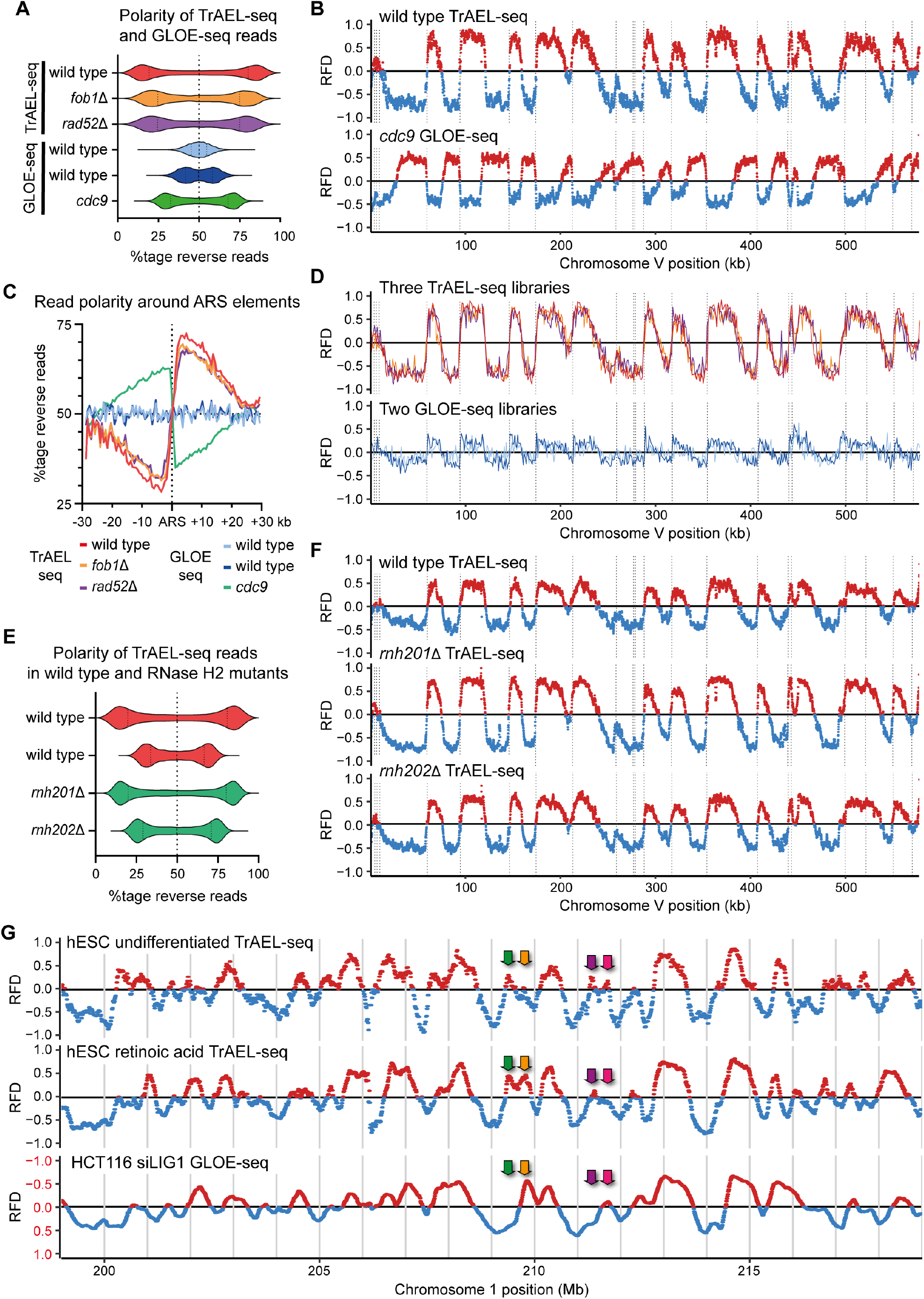
TrAEL-seq is highly sensitive to replication fork direction. **A**: Polarity of TrAEL-seq and GLOE-seq reads assessed in 1kb windows across the genome excluding windows overlapping multi-copy regions, presented as the percentage of total reads that map to the reverse strand. The dotted line marks 50%, which equates to an absence of strand bias. GLOE-seq wild-type samples (SRA accessions: SRX6436839 and SRX6436840) were derived from asynchronous log phase cells growing in YPD, as were all the TrAEL-seq samples. The *cdc9* dataset is of synchronised cells depleted of the DNA ligase Cdc9 (SRA accession: SRX6436838). **B**: RFD plots for TrAEL-seq wild type and GLOE-seq Cdc9 depletion data (SRA accession: SRX6436838) across chromosome V, calculated as (R-F)/(R+F) where R and F indicate reverse and forward reads respectively. RFD was calculated for 1000 bp sliding windows spaced every 100 bp. Vertical dotted lines show locations of ARS elements. **C**: Average read polarity of TrAEL-seq and GLOE-seq datasets across 30 kb windows either side of annotated ARS elements. Calculated as the %tage of reverse reads amongst all reads. **D**: Signal strength and reproducibility amongst TrAEL-seq and GLOE-seq datasets. RFD was calculated in 1000 bp windows spaced every 1000 bp and shown as continuous lines. Wild-type (red), *fob1*Δ (orange) and *rad52*Δ (purple) datasets are plotted on the upper graph and show indistinguishable replication profiles. Two wild type GLOE-seq datasets are overlaid on the lower graph (SRA accessions: SRX6436839 and SRX6436840). TrAEL-seq and GLOE-seq datasets all derive from asynchronous cultures harvested during log phase growth in YPD [36]. Vertical dotted lines show locations of ARS elements. **E**: Read polarity plot as in A for two replicate wild type TrAEL-seq datasets compared to the RNase H2 mutants *rnh201*Δ and *rnh202*Δ. **F**: RFD plots as in B for *rnh201*Δ and *rnh202*Δ mutants with matched wild-type control. **G:** RFD plots of TrAEL-seq data for asynchronous undifferentiated and retinoic acid treated hESCs, compared to GLOE-seq data for LIG1-depleted HCT116 cells (average of SRA accessions: SRX7704535 and SRX7704534). RFD was calculated in 250 kb sliding windows spaced every 10 kb. Note that the polarity of the HCT116 data has been inverted to aid comparison with TrAEL-seq samples, this is highlighted by the scale being labelled in red.

A replication signal of the same polarity was noted in wild-type samples profiled by GLOE-seq but only very weakly, whereas all three TrAEL-seq libraries display very strong read polarity differences and essentially identical replication profiles (Fig. 3A and D) [36]. The only major difference is observed in the rDNA, which is highly polarised towards reverse forks in wild-type and *rad52*Δ due to the RFB asserting replication directionality whereas this polarisation is absent in *fob1*Δ cells (Fig. S3B). As Sriramachandran *et al*. noted for GLOE-seq [36], the read polarity of this replication signal is opposite to what would be expected from labelling of 3’ ends in normal forks. There should never be less 3’ ends on the lagging strand than the leading strand, yet up to 90% of TrAEL-seq reads emanate from the leading strand. To explain the GLOE-seq signal, Sriramachandran *et al*. suggested that GLOE-seq labels sites at which DNA is nicked during removal of mis-incorporated ribonucleotides [36]. To test this idea, we generated TrAEL-seq libraries from *rnh201*Δ and *rnh202*Δ mutants that lack key components of RNase H2, the main enzyme that cleaves DNA at mis-incorporated ribonucleotides, along with a wild-type control [68, 69]. Strikingly, both read polarity bias and RFD plots for these mutants are equivalent to wild-type, showing that the leading strand bias of TrAEL-seq reads is not caused by RNase H2 and therefore is unlikely to arise through excision of mis-incorporated ribonucleotides (Fig. 3E, F). One further observation in this regard is that END-seq data also shows a polarity bias, albeit weak, that parallels the polarity bias in TrAEL-seq data generated from the same cells (Fig. S3C), suggesting that double stranded ends are also formed during normal replication (see Discussion).

We then asked if an equivalent bias is observed in the hESC libraries. The limited read coverage in these libraries only allowed read polarity to be determined in 250 kb windows, but nonetheless a striking variation was observed across the genome in RFD plots (Fig. 3G). Importantly, these profiles were very similar between undifferentiated and differentiated hESCs and cannot therefore simply result from noise; this can be observed in example RFD plots, but is also clear in a scatter plot which shows that the average read polarity within each window correlates between the datasets with R^2=0.73 (Fig. S3D). Furthermore, comparison to GLOE-seq results from a LIG1-depleted human cell line that is defective in Okazaki fragment ligation again revealed a striking similarity to the hESC TrAEL-seq data by RFD plot and scatter plot, although with the opposite polarity as expected (Fig. 3F and S3D) [36]. Interestingly, a subset of origins were not active in one condition or cell type, underlining the fact that replication origin usage is more variable in mammals than in yeast and changes, for example, during stem cell differentiation (Fig. 3G, example origins are indicated by coloured arrows) [70].

We therefore conclude that TrAEL-seq unexpectedly provides remarkably accurate replication profiles. These profiles are highly reproducible and can be obtained from wild-type cells without need for cell synchronisation, sorting or labelling.

### Environmental impacts on replication timing and fork progression

Finally, we asked whether TrAEL-seq can reveal replication changes or DNA damage, and in particular whether we can detect collisions between transcription and replication machineries.

Since all the yeast libraries generated up to this point had yielded essentially identical DNA replication profiles outside the rDNA, we were first keen to ensure that changes in replication profile are indeed detectable. We therefore examined cells lacking Clb5, a yeast cyclin B that plays a key role in the activation of late firing replication forks [71]. The TrAEL-seq profile of *clb5*Δ was very similar to wild-type across most of the genome, but certain origins were clearly absent or strongly repressed, resulting in extended tracts of DNA synthesis from adjacent origins (Fig. 4A, green arrows). This is as predicted for *clb5*Δ mutants and confirms that TrAEL-seq is indeed sensitive to changes in replication profile.

**Figure 4:**
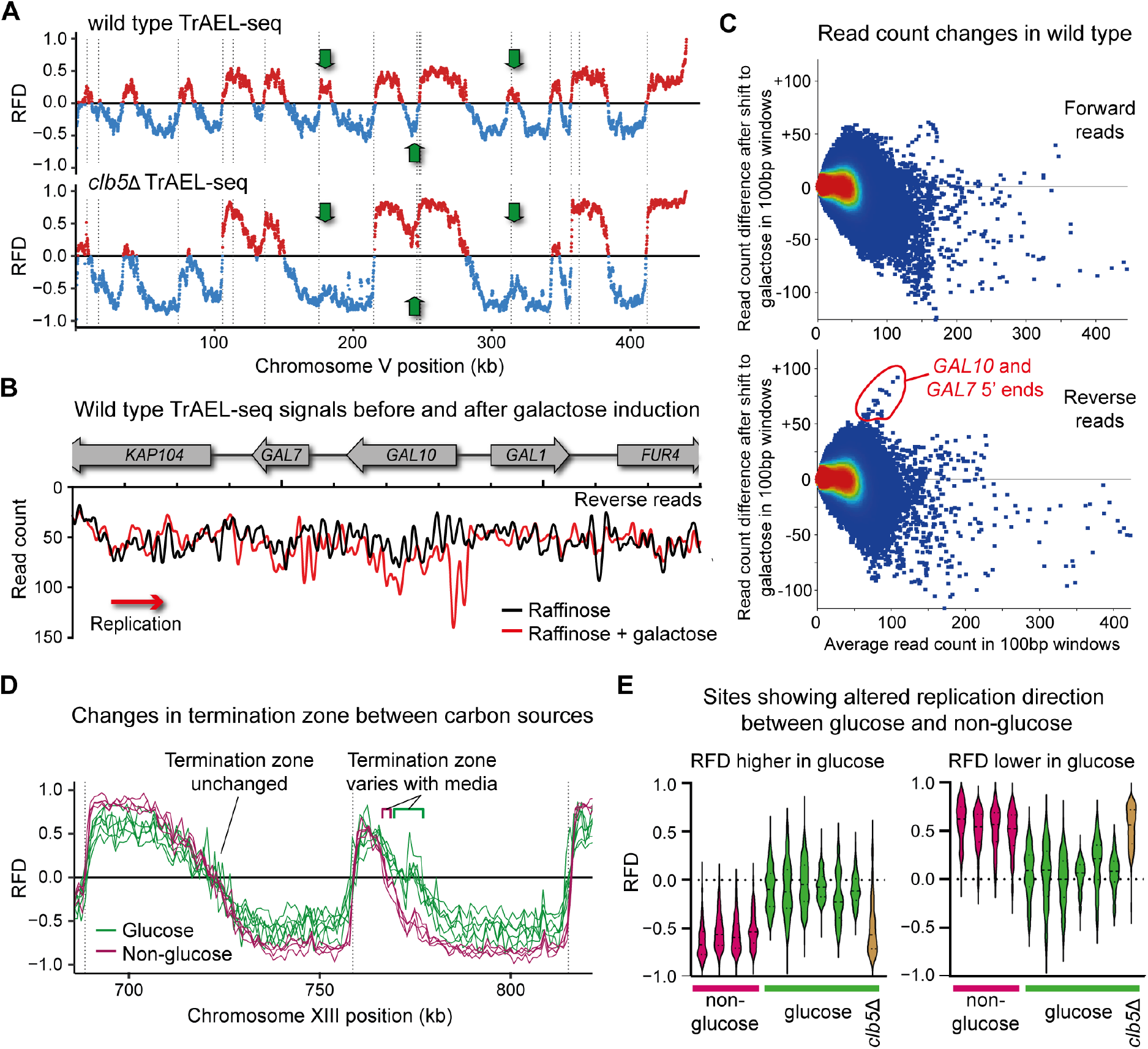
Detection of replication variation using TrAEL-seq. **A**: RFD plot for *clb5*Δ versus wild type. Arrows indicate ARS elements that are not activated in the absence of Clb5. **B**: Line plot showing reverse strand TrAEL-seq read counts across the GAL genes for wild-type cells maintained on YP raffinose or 5 hours after addition of galactose to 2%. Reads were quantified in 100 bp sliding windows spaced every 10 bp. **C**: MA plots showing the change in read count against the average read count for each 100 bp window in the single copy genome between cells maintained on raffinose and cells exposed to galactose. Separate plots are shown for forward and reverse reads. **D**: Example location in which a termination zone differs depending on carbon source. RFDs were calculated in 1 kb windows spaced every 1 kb. Green lines show cells grown on glucose and purple lines cells grown on raffinose or raffinose plus galactose. **E:** Violin plots of regions showing large and significant RFD differences between cells grown on glucose (green) and those growing on raffinose or raffinose plus galactose (purple). RFDs were calculated across the single copy genome in 1 kb windows, then each window compared between the two sets by *t*-test with a Benjamini and Hochberg correction. Windows were then further filtered for those with a difference in RFD>0.4 between the two sets, leaving a set of 290 out of 12182 (2.3%). Plots were split based on the direction of the difference in RFD for clarity. Individual violin plots represent respectively: wild type on raffinose, wild type on raffinose + galactose, *dnl4*Δ *rad51*Δ on raffinose, *dnl4*Δ *rad51*Δ on raffinose + galactose, then *fob1*Δ, wild type, *rad52*Δ, wild type, *rnh201*Δ, *rnh202*Δ, *clb5*Δ all on glucose.

We then engineered collisions between RNA polymerase II and the replisome by changing growth conditions to strongly induce certain genes; specifically, we added galactose to cells growing on raffinose, which strongly induces expression of galactose metabolising genes including *GAL1, GAL7* and *GAL10*. Although these genes are adjacent, *GAL1* is transcribed co-directionally with the replication fork whereas *GAL7* and *GAL10* are orientated head-on to the fork. On one hand, stalled replication forks have not been observed at this locus by 2D gels [65], but conversely the strong activation of the *GAL1-10* promoter has proved highly recombinogenic in various assays [72–74]. We performed these experiments in wild-type cells and in a strain lacking both Dnl4, the DNA ligase required for non-homologous end joining, and Rad51, the recA ortholog which mediates strand invasion for homologous recombination. *dnl4*Δ *rad51*Δ double mutants should be unable to repair DSBs irrespective of cell cycle phase and therefore should accumulate DSBs.

Analysis of TrAEL-seq read densities across the *GAL* gene cluster provided evidence for transcription-associated replication fork stalling but not in the expected location. Peaks of reverse reads formed at the 5’ end of the *GAL10* gene, and also of the *GAL7* gene although the latter was less prominent, which suggests that the replication fork is stalled by chromatin or proteins bound at the promoter (Fig. 4B and S4A). The read accumulation is not dramatic, but compared to the rest of the single copy genome these sites showed the largest increase in read count between cells on raffinose only and those on raffinose plus galactose (Fig. 4C and S4B). In contrast, we observed only slight evidence of reads accumulating in the *GAL10* or *GAL7* open reading frames that would indicate collisions between the replication and transcription machineries (Fig. 4B and S4A), nor did we detect a major difference between *dnl4*Δ *rad51*Δ and the wild type showing that promoter signals must represent fork stalling events that are rarely processed to recombinogenic DSBs (Fig. S4A and S4B). Furthermore, the region in which replication forks passing through the *GAL* locus encounter oncoming forks from ARS211 was unchanged on galactose meaning that delays caused by fork stalling must be very transient (Fig. S4C).

Unexpectedly, we noted changes in termination zones elsewhere in the genome when comparing the four samples from this experiment, which were grown on raffinose or raffinose with galactose, to other wild-type and mutant TrAEL-seq libraries for which cells were grown on glucose (see for example Fig. 4D). Comparing cells based on growth media rather than genotype, we discovered significant and substantial (p<0.01, average RFD change >0.4) differences in RFD for ^~^2% of the single copy genome (Fig. 4E). The most prominent differences affected a subset of termination zones where the average site at which forks converge moved by up to 10 kb. This change would be most easily attributed to a change in replication timing, and indeed the *clb5*Δ mutant, although grown on glucose, showed the same average RFD at the media-dependent sites as the cells grown on raffinose or raffinose with galactose (Fig. 4E). This suggests that the timing of replication firing is altered depending on carbon source, consistent with a previous report that Clb5 nuclear import is suppressed in yeast growing in ethanol [75].

Together, these data show that replication profiling by TrAEL-seq is sufficiently sensitive to reveal subtle differences in fork direction and processivity.

## Discussion

Here we have demonstrated that TrAEL-seq maps resected DSBs, sites of replication fork stalling and normal DNA replication patterns genome-wide and with base-pair resolution. Methods to map the 3’ ends of resected DNA are desirable for genome-wide studies of homologous recombination as these are the critical species that undergo strand invasion. Similarly, detection of DNA 3’ ends at stalled replication forks is an important indicator of potentially recombinogenic intermediates. TrAEL-seq profiles all these species with excellent signal-to-noise and therefore provides a general method for the detection of DNA processing events that could result in genome instability. It is interesting to note that the primary source of noise in TrAEL-seq is actually normal replication forks. This raises questions as to the frequency with which leading strand 3’ ends become detached during normal replication (discussed below), but also provides a major unanticipated application for the method. In contrast to other methods for profiling replication fork directionality (notably through Okazaki fragment sequencing), TrAEL-seq works in wild-type cells, requires neither labelling nor synchronisation of cells and does not involve complex sample preparation procedures, making TrAEL-seq versatile and straightforward to implement across a range of experimental contexts.

### A proposed mechanism for replication fork detection by TrAEL-seq

TrAEL-seq was designed to detect free 3’ ends of single stranded DNA and was not expected to label undisturbed replication forks in normal cells. Why therefore is TrAEL-seq so sensitive to replication fork direction? The mechanism responsible must frequently rearrange replication forks to make the leading strand 3’ end accessible to TdT while the lagging strand 3’ end remains largely inaccessible. We suggest that transient fork reversal would have this effect, yielding TdT-accessible leading strand ends without irreversible changes in fork structure (Fig. 5, free 3’ ends labelled with green dots). This mechanism would require surprisingly frequent fork reversal, but for TrAEL-seq labelling the reversal required is minimal - in reality only a flap displacement (Fig. 5, left and middle structures). Although DNA replication is highly processive overall, *in vitro* measurements have shown that the yeast leading and lagging strand polymerases dissociate after less than 1 kb of DNA synthesis [76], and we suggest this provides sufficient opportunity for helicases to access and unwind the nascent leading strand. A small subset of these events likely result in sufficient reversal for the nascent lagging and leading strands to anneal and form the replication-linked double-stranded DNA ends that we detect by END-seq (Fig. 5, middle and right structures and Fig. S3C).

**Figure 5:**
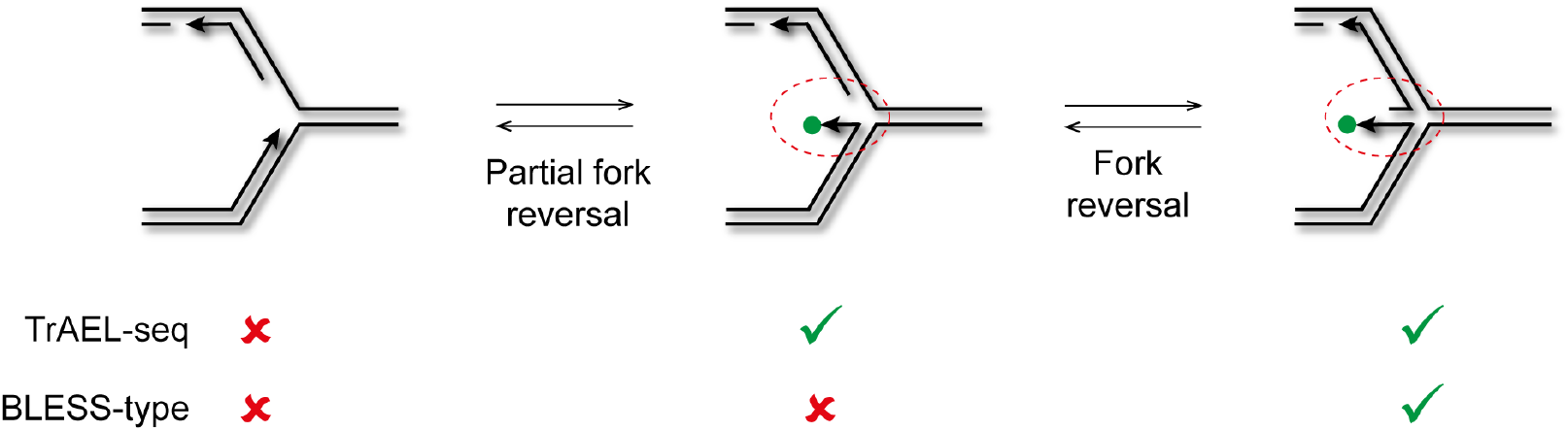
Proposed mechanism for replication fork detection by TrAEL-seq. Replication forks that would normally be undetectable by TrAEL-seq undergo very limited reversal to yield a free 3’ end that can be labelled by TdT (green dot, middle structure). Further reversal yields a double stranded end that can be labelled by TrAEL-seq or BLESS-type methods. Red circles highlight the area of difference between the structures.

Alternatively, it is possible that the TrAEL-seq replication signal derives from cleaved replication forks but we think this highly unlikely for the following reasons: 1) the *rad52*Δ mutant used here had almost no growth defect and showed no detectable difference in TrAEL-seq profile, and 2) there is no difference in detection of early and late replicating genome regions in TrAEL-seq, whereas the activity of structure specific endonucleases that could cleave replication forks is tightly restricted to G2/M [77]. Replication-linked double-stranded DNA ends have been clearly observed by BLESS-type methods in cells exposed to replication stress [13, 15, 78] and interpreted as evidence that replication forks are cleaved either during the restart process or as a pathogenic end point. However, fork cleavage is not required to initiate recombination during replication fork restart [79], and it is quite possible that apparent DSBs are actually double stranded ends of reversed forks. Direct observation of cleaved forks at the rDNA RFB has been reported based on southern blot [47, 48, 53], but we note that these signals could also arise from fork reversal (Fig. S5). This distinction is important as cleaved forks must be resolved by recombination of some sort whereas reversed forks can revert by Holliday Junction migration. Overall, the existence of frequent DSBs in wild-type cells under normal conditions (quantified at 1 DSB per cell per S-phase for the RFB alone [13]) is hard to reconcile with the minimal growth phenotype of mutants lacking critical DNA repair factors such as Rad52, and we suggest that the vast majority of such events detected by TrAEL-seq and other DNA end-mapping methods are actually reversed replication forks that are rapidly resolved by fork migration.

### Complementary methods probe different aspects of DNA damage

Although TrAEL-seq and the recently described GLOE-seq methodology in theory act equivalently by labelling and profiling DNA 3’ ends, we find that these methods have completely different strengths and weaknesses. TrAEL-seq proves superior for detection of replication fork direction and stalling, which likely arises through a sensitivity to replication fork structure. In contrast, the DNA denaturing step required for GLOE-seq labelling erases fork structure and reveals real accumulations of strand breaks as opposed to conformational changes in the replication fork. Therefore future studies employing both methods in parallel are likely to be particularly informative for understanding the dynamics of replication forks on encountering obstacles. It should also be noted that the lack of a denaturing step in TrAEL-seq makes it insensitive to single-strand breaks and nicks, and therefore GLOE-seq is much better suited for detection of such ends.

Genome-wide analysis of DNA processing events requires high-resolution methods that can detect changes at both 5’ and 3’ DNA ends. BLESS-type methods degrade or fill-in 3’ ends to yield the location of matching 5’ ends, and our implementation of TrAEL-seq now provides a complementary method to map 3’ ends. We suggest that for dissecting mechanisms of DSB processing and repair, these methods will be most powerful when employed together. In addition to the TrAEL-seq protocol, we therefore also provide an implementation of BLESS / END-seq that utilises small numbers of cells and follows the same library construction procedure as TrAEL-seq, making processing of the same sample in parallel by both methods straightforward. Indeed, we have successfully performed TrAEL-seq and END-seq on two halves of the same agarose plug containing 10 million cells.

For general replication analysis, existing methods profile either fork direction or origin timing whereas acquisition of information on both parameters from the same samples would be very helpful. The ability of TrAEL-seq to obtain replication direction profiles from asynchronous unlabelled wild type cells will allow easy integration with other methods. For example, ethanol fixed cells collected for sort-seq [24] could also be profiled by TrAEL-seq to provide both replication timing and direction. However, some adjustments will be needed when combining TrAEL-seq with replication timing methods that involve labelling with deoxyuridine derivatives (e.g.: REPLI-seq) as USER is employed in TrAEL-seq to elute libraries prior to amplification.

Overall, TrAEL-seq provides a unique addition to complement existing methods for genome-wide analysis of DNA replication and DNA damage. The relatively simple experimental protocol, high signal-to-noise ratio and lack of requirement for treatment or purification of cells prior to harvest should render TrAEL-seq particularly suitable for a wide range of experimental systems.

## Materials and Methods

### Yeast strains and culture

Strains used are listed in Table S1. All media components were purchased from Formedium, all media was filter sterilised. YP media was supplemented with the given carbon source from 20% filter-sterilised stock solutions. For growth to log phase, cells were inoculated in 4 ml media and grown for ^~^6 h at 30°C with shaking at 200 rpm before dilution at ^~^1:10,000 in 25 ml YPD (1:500 for YP raffinose or 1:2000 for synthetic complete media) and growth continued at 30°C 200 rpm for ^~^18 hours until OD reached 0.4-0.7 (mid-log). Cells were centrifuged 1 min at 4,600 rpm, resuspended in 70% ethanol at 1×10^7^ cells/ml and stored at −70°C. For meiosis, SK1 *dmc1*Δ diploid cells from a glycerol stock were patched overnight on YP 2% Glycerol then again for 7 hours on YP 4% glucose before inoculating in 4 ml YPD and growth for 24 hours, then inoculated to OD 0.2 in 20 ml YP acetate for overnight growth to ^~^4×10^7^ cells/ml in a 100 ml flask at 30°C with shaking at 200 rpm. Meiosis was initiated by washing cells once with 20ml SPO media (0.3% KOAc, 5 mg/L uracil, 5 mg/L histidine, 25 mg/L leucine, 12.5 mg/L tryptophan, 0.02% raffinose) then resuspending in 20 ml SPO media and incubating for 7 hours at 30°C in a 100 ml flask with shaking at 250 rpm. Cells were harvested and fixed with 70% ethanol as above.

### Culture and differentiation of hESCs

Undifferentiated H9 hESCs were maintained on Vitronectin-coated plates (ThermoFisher Scientific) in TeSR-E8 media (StemCell Technologies). To induce differentiation, hESCs were grown in N2B27 media supplemented with 3 μM retinoic acid for 72 hours. N2B27 media consists of a 1:1 mixture of DMEM/F12 and Neurobasal, 0.5x N2 supplement, 0.5x B27 supplement, 1x nonessential amino acids, 2mM L-Glutamine, 1x Penicillin/Streptomycin (all from ThermoFisher Scientific) and 0.1 mM β-mercaptoethanol. All hESCs were cultured in 5% O_2_, 5% CO_2_ at 37°C.

### Agarose embedding of yeast cells

Cells in ethanol (1-3×10^7^ per plug) were pelleted in round bottom 2 ml tubes by centrifuging 30 s 20,000 g, washed once in 1 ml PFGE wash buffer (10 mM Tris HCl pH 7.5, 50 mM EDTA) and resuspended in 60 μl same with 1 μl lyticase (17 U/ μl in 10 mM KPO_4_ pH7, 50% glycerol, Merck >2000 U/mg L2524). Samples were heated to 50°C for 1-10 min before addition of 40 μl molten CleanCut agarose (Bio-Rad 1703594), vortexing vigorously for 5 s before pipetting in plug mould (Bio-Rad 1703713) and solidifying 15-30 min at 4°C. Each plug was transferred to a 2 ml tube containing 500 μl PFGE wash buffer with 10 μl 17 U/μl lyticase and incubated 1 hour at 37°C. Solution was replaced with 500 μl PK buffer (100 mM EDTA pH 8, 0.2% sodium deoxycholate, 1% sodium N-lauroyl sarcosine, 1 mg/ml Proteinase K) and incubated overnight at 50°C. Plugs were rinsed with 1 ml TE, then washed three times with 1 ml TE for 1-2 hours at room temperature with rocking, 10 mM PMSF was added to the second and third washes from 100 mM stock (Merck 93482). Plugs were then digested 1 h at 37°C with 1 μl 1000 U/ml RNase T1 (Thermo EN0541) in 200 μl TE. RNase A was not used as it binds strongly to single stranded DNA [80]. Plugs were stored in 1 ml TE at 4°C and are stable for >1 year.

### Agarose embedding of hESC cells

Cells were detached using Accutase, counted and 1×10^6^ cells were washed once in 5 ml L buffer (10 mM Tris HCl pH 7.5, 100 mM EDTA, 20 mM NaCl) and resuspended in 60 μl L buffer in a 2 ml tube. Samples were heated to 50°C for 2-3 min before addition of 40 μl molten CleanCut agarose (Bio-Rad 1703594), vortexing vigorously for 5 s before pipetting in plug mould (Bio-Rad 1703713) and solidifying 15-30 min at 4°C. Each plug was transferred to a 2 ml tube containing 500 μl digestion buffer (10 mM Tris HCl pH 7.5, 100 mM EDTA, 20 mM NaCl, 1% sodium N-lauroyl sarcosine, 0.1 mg/ml Proteinase K) and incubated overnight at 50°. Plugs were washed and RNase T1 treated as for yeast.

### TrAEL-seq library preparation and sequencing

Please note that a detailed TrAEL-seq protocol is provided in Supplementary Material, and up-to-date protocols are available from the Houseley lab website https://www.babraham.ac.uk/our-research/epigenetics/jon-houseley/protocols

Preparation of TrAEL-seq adaptor 1: DNA oligonucleotide was synthesised and PAGE purified by Sigma-Genosys (Merck):

[Phos]NNNNNNNNAGATCGGAAGAGCGTCGTGTAGGGAAAGAGTGTUGCGCAGGCCATTGGCC[BtndT]GCGCUACACTCTTTCCCTACACGACGCT

This oligonucleotide was adenylated using the 5’ DNA adenylation kit (NEB, E2610S) as follows: 500 pMol DNA oligonucleotide, 5 μl 10x 5’ DNA Adenylation Reaction Buffer, 5 μl 1 mM ATP, 5 μl Mth RNA Ligase in a total volume of 50 μl was incubated for 1 h at 65°C then 5 min at 85°C. Reaction was extracted with phenol:chloroform pH 8, then ethanol precipitated with 10 μl 3M NaOAc, 1 μl GlycoBlue (Thermo AM9515), 330 μl ethanol and resuspended in 50 μl 0.1x TE.

Preparation of TrAEL-seq adaptor 2: DNA oligonucleotide was synthesised and PAGE purified by Sigma-Genosys (Merck):

[Phos]GATCGGAAGAGCACACGTCTGAACTCCAGTCUUUUGACTGGAGTTCAGACGTGTGCTCTTCCGATC*T

Oligonucleotide was annealed before use: 20 μl 100 pM/μl oligonucleotide and 20 μl 10x T4 DNA ligase buffer (NEB) in 200 μl final volume were incubated in a heating block 95°C 5 min then block was removed from heat and left to cool to room temperature over ^~^2 h.

Sample preparation: ½ an agarose plug was used for each library (cut with a razor blade), hereafter referred to as a plug for simplicity. All incubations were performed in 2 ml round bottomed tubes (plugs break easily in 1.5 ml tubes), or 15 ml tubes for high volume washes. For *Sfi*I digestion, a plug was equilibrated 30 min in 100 μl 1x CutSmart buffer (NEB), digested overnight at 50°C with 1 μl 20 U/μl *Sfi*I (NEB R0123S) in 200 μl 1x CutSmart buffer, and then rinsed with 1xTE.

Tailing and ligation: Plugs were equilibrated once in 100 μl 1x TdT buffer (NEB) for 30 min at room temperature, then incubated for 2 h at 37°C in 100 μl 1x TdT buffer containing 4 μl 10 mM ATP and 2 μl Terminal Transferase (NEB M0315L). Plugs were rinsed with 1 ml tris buffer (10 mM Tris HCl pH 8.0), equilibrated in 100 μl 1x T4 RNA ligase buffer (NEB) containing 40 μl 50% PEG 8000 for 1 hour at room temperature then incubated overnight at 25°C in 100 μl 1x T4 RNA ligase buffer (NEB) containing 40 μl 50% PEG 8000, 1 μl 10 pM/μl TrAEL-seq adaptor 1 and 1 μl T4 RNA ligase 2 truncated KQ (NEB M0373L). Plugs were then rinsed with 1 ml tris buffer, transferred to 15 ml tubes and washed three times in 10 ml tris buffer with rocking at room temperature for 1-2 hours each, then washed again overnight under the same conditions.

DNA processing: Plugs were equilibrated for 15 min with 1 ml agarase buffer (10 mM Bis-Tris-HCl, 1 mM EDTA pH 6.5), then the supernatant removed and 50 μl agarase buffer added. Plugs were melted for 20 min at 65°C, transferred for 5 min to a heating block pre-heated to 42°C, 1 μl β-agarase (NEB M0392S) was added and mixed by flicking without allowing sample to cool, and incubation continued at 42°C for 1 h. DNA was ethanol precipitated with 25 μl 10 M NH_4_OAc, 1 μl GlycoBlue, 330 μl of ethanol and resuspended in 10 μl 0.1xTE. 40 μl reaction mix containing 5 μl Isothermal amplification buffer (NEB), 3 μl 100 mM MgSO_4_, 2μl 10 mM dNTPs and 1μl Bst 2 WarmStart DNA polymerase (NEB M0538S) was added and sample incubated 30 min at 65°C before precipitation with 12.5 μl 10 M NH_4_OAc, 1 μl GlycoBlue, 160 μl ethanol and re-dissolving pellet in 130 μl 1xTE. The DNA was transferred to an AFA microTUBE (Covaris 520045) and fragmented in a Covaris E220 using duty factor 10, PIP 175, Cycles 200, Temp 11°C, then transferred to a 1.5 ml tube containing 8 μl pre-washed Dynabeads MyOne streptavidin C1 beads (Thermo, 65001) re-suspended in 300 μl 2xTN (10 mM Tris pH 8, 2 M NaCl) along with 170 μl water (total volume 600 μl) and incubated 30 min at room temperature on a rotating wheel. Beads were washed once with 500 μl 5 mM Tris pH 8, 0.5 mM EDTA, 1 M NaCl, 5 min on wheel and once with 500 μl 0.1x TE, 5 min on wheel before re-suspension in 25 μl 0.1x TE.

Library preparation: TrAEL-seq adaptor 2 was added using a modified NEBNext Ultra II DNA kit (NEB E7645S): 3.5 μl NEBNext Ultra II End Prep buffer, 1 μl 1 ng/μl sonicated salmon sperm DNA (this is used as a carrier) and 1.5 μl NEBNext Ultra II End Prep enzyme were added and reaction incubated 30 min at room temperature and 30 min at 65°C. After cooling, 1.25 μl 10 pM/μl TrAEL-seq adaptor 2, 0.5 μl NEBNext ligation enhancer and 15 μl NEBNext Ultra II ligation mix were added and incubated 30 min at room temperature. The reaction mix was removed and discarded and beads were rinsed with 500 μl wash buffer (5 mM Tris pH 8, 0.5 mM EDTA, 1 M NaCl) then washed twice with 1 ml wash buffer for 10 min on wheel at room temperature and once for 10 min with 1 ml 0.1x TE. Libraries were eluted from beads with 11 μl 1x TE and 1.5 μl USER enzyme (NEB) for 15 min at 37°C, then again with 10.5 μl 1x TE and 1.5 μl USER enzyme (NEB) for 15 min at 37°, and the two eluates combined.

Library amplification: Amplification was performed with components of the NEBNext Ultra II DNA kit (NEB E7645S) and a NEBNext Multiplex Oligos set (e.g. NEB E7335S). An initial test amplification was used to determine the optimal cycle number for each library. For this, 1.25 μl library was amplified in 10 μl total volume with 0.4 μl each of the NEBNext Universal and any NEBNext Index primers with 5 μl NEBNext Ultra II Q5 PCR master mix. Cycling program: 98°C 30s then 18 cycles of (98°C 10 s, 65°C 75 s), 65°C 5 min. Test PCR was cleaned with 8 μl AMPure XP beads (Beckman A63881) and eluted with 2.5 μl 0.1x TE, of which 1 μl was examines on a Bioanalyser high sensitivity DNA chip (Agilent 5067-4626). Ideal cycle number should bring final library to final concentration of 1-3 nM, noting that the final library will be 2-3 cycles more concentrated than the test anyway. 21 μl of library was then amplified with 2 μl each of NEBNext Universal and chosen Index primer and 25 μl NEBNext Ultra II Q5 PCR master mix using same conditions as above for calculated cycle number. Amplified library was cleaned with 40 μl AMPure XP beads (Beckman A63881) and eluted with 26 μl 0.1x TE, then 25 μl of this was again purified with 20μl AMPure XP beads and eluted with 11μl 0.1x TE. Final libraries were quality controlled and quantified by Bioanalyser (Agilent 5067-4626) and KAPA qPCR (Roche KK4835).

Libraries were sequenced either on an Illumina MiSeq as 50bp Single Read or an Illumina NextSeq 500 as High Output 75 bp Single End by the Babraham Institute Next Generation Sequencing facility.

### END-seq library preparation

Note: This protocol is based on the original described by Canela *et al*. [11] but has a critical difference: the exonuclease-mediated blunting step designed for topoisomerase II ends did not work well on the two test substrates we use in yeast genomic DNA. Instead, best results were obtained by blunting 2 hours or over-night with Klenow, which outperformed T4 DNA polymerase or a commercial DNA blunting kit.

Preparation of END-seq adaptor 1: DNA oligonucleotide was synthesised and PAGE purified by Sigma-Genosys (Merck), sequence is as described by Canela *et al*. [11]:

[Phos]GATCGGAAGAGCGTCGTGTAGGGAAAGAGTGUU[BtndT]U[BtndT]UUACACTCTTTCCCTACACGACGCTCTTCCGATC*T

Annealed as for TrAEL-seq adaptor 2 above.

Preparation of END-seq adaptor 2c: DNA oligonucleotide was synthesised and PAGE purified by Sigma-Genosys (Merck), modified from Canela *et al*. [11] to prevent homo-dimers of adaptor from amplifying:

[Phos]GATCGGAAGAGCTATTATTTAAATTTTAATTUGACTGGAGTTCAGACGTGTGCTCTTCCGATC*T

Annealed as for TrAEL-seq adaptor 2 above.

Sample preparation: ½ an agarose plug was used for each library (cut with a razor blade), hereafter referred to as a plug for simplicity. All incubations were performed in 2 ml round bottomed tubes (plugs break easily in 1.5 ml tubes), or 15 ml tubes for high volume washes. For *Sfi*I digestion, plug was equilibrated 30 min in 100 μl CutSmart buffer (NEB), digested overnight at 50°C with 1 μl 20 U/μl *Sfi*I (NEB R0123S) then rinsed with 1xTE.

Blunting and ligation: The plug was equilibrated for 1 h at room temperature in 100 μl NEBuffer 2 with 0.1 mM dNTPs, then blunted overnight at 37°C in 100 μl NEBuffer 2 with 0.1 mM dNTPs and 1 μl Klenow (NEB M0210S). After rinsing twice with 1 ml tris buffer, plug was transferred to a 15 ml tube and washed three times for 15 min each with 10 ml tris buffer on rocker at room temperature before transfer to a new 2 ml tube. The plug was equilibrated with 100 μl CutSmart buffer containing 5 mM DTT and 1 mM dATP for 1 hour at room temperature before incubation for 2 h at 37°C in another 100 μl of the same buffer containing 1 μl Klenow exo-(NEB M0212S) and 1 μl T4 PNK (NEB M0201S). Plug was rinsed twice with 1 ml tris buffer then washed once with 10 ml of tris buffer for 15 min as above then returned to a 2 ml tube. The plug was equilibrated for 1 h at room temperature in 100 μl 1x Quick Ligation buffer (NEB B6058S) containing 2.7 μl END-seq adaptor 1, then overnight at 25°C with another 100 μl of the same buffer containing 2.7 μl END-seq adaptor 1 and 1 μl high concentration T4 DNA Ligase (NEB M0202M). After rinsing twice with 1 ml tris buffer, plug was transferred to a 15 ml tube and washed three times for 1-2 h each with 10 ml tris buffer on rocker at room temperature, then again overnight.

DNA purification and library construction: The plug was transferred to a 1.5 ml tube and equilibrated 15 min with 1 ml agarase buffer (10 mM Bis-Tris-HCl, 1 mM EDTA pH 6.5), then the supernatant removed and 50 μl agarase buffer added to the plug. Plug was melted 20 min at 65°C then transferred for 5 min to a heating block pre-heated to 42°C, 1 μl beta-agarase (NEB M0392S) was added and mixed by flicking without allowing sample to cool, and incubation continued at 42°C for 1 h. DNA was ethanol precipitated with 25 μl 10 M NH_4_OAc, 1 μl GlycoBlue, 330 μl of ethanol and resuspended in 130 μl 1x TE, 15 min at 65°C. From here, samples were sonicated, purified and library construction performed as for TrAEL-seq, except that END-seq adaptor 2c was substituted for TrAEL-seq adaptor 2.

### in vitro *TrAEL activity and qPCR assays*

For *in vitro* assay, 0.5 μl 10 mM DNA oligonucleotide CGCGGTAATTCCAGCTCCAA was treated with or without 0.5 μl TdT in 20 μl 1x TdT buffer containing 0.8 μl 10 mM ATP for 30 min at 37°C. Reactions were purified by phenol:chloroform extraction and ethanol precipitation, and resuspended in 5 μl 10 mM Tris pH8. This was ligated to 1 μl TrAEL-seq adaptor 1 in 20 μl 1x T4 RNA ligase buffer containing 8 μl 50% PEG 8000 and 1 μl T4 RNA ligase 2 truncated KQ overnight at 25°C. Reactions were resolved on a 15% PAGE / 8M urea gel and stained with SYBR Gold (Thermo S11494) as per manufacturer’s instructions.

For qPCR, DNA taken at the indicated points was diluted with water and qPCR reactions performed in 10 μl volumes with 5 μl Maxima SYBR Green qPCR Master Mix (Thermo K0222) and 0.3 μl each 10 μM primer dilution, primers are listed in Table S2. Cycling parameters: 95°C 10min, 40x (95°C 15 s, 70°C 60 s) on a Bio-Rad CFX-96.

### Data analysis

Unique Molecular Identifier (UMI) deduplication and mapping: Scripts used for UMI-handling as well as more detailed information on the processing are available here: https://github.com/FelixKrueger/TrAEL-seq). Briefly, TrAEL-seq reads are supposed to carry an 8 bp in-line barcode (UMI) at the 5’-end, followed by a variable number of 1-3 thymines (T). The script TrAELseq_preprocessing.py removes the first 8bp (UMI) of a read and adds the UMI sequence to the end of the readID. After this, up to 3 T (inclusive) at the start of the sequence are removed. Following this UMI and Poly-T pre-processing, reads underwent adapter- and quality trimming using Trim Galore (v0.6.5; default parameters; https://github.com/FelixKrueger/TrimGalore). UMI-pre-processed and adapter-/quality trimmed files were then aligned to the respective genome using Bowtie2 (v2.4.1; option: --local; http://bowtie-bio.sourceforge.net/bowtie2/index.shtml) using local alignments. Finally, alignment results files were deduplicated using UmiBam (v0.2.0; https://github.com/FelixKrueger/Umi-Grinder). This procedure deduplicates alignments based on the mapping position, read orientation as well as the UMI sequence.

UMI de-duplicated mapped reads were imported into SeqMonk v1.47 (https://www.bioinformatics.babraham.ac.uk/projects/seqmonk/) and immediately truncated to 1 nucleotide at the 5’ end, representing the last nucleotide 5’ of the strand break. Reads were then summed in running windows or around features as described in figure legends. Windows overlapping with non-single copy regions of the genome were filtered (rDNA, 2μ, mtDNA, *CUP1*, sub-telomeric regions, Ty elements and LTRs), and total read counts across all included windows were normalised to be equal. Scatter plots and average profile plots were generated in SeqMonk, and in the latter case the data was exported and plots re-drawn in GraphPad Prism 8.

For read count quantification and RFD plots, data was first imported into SeqMonk v1.47 and truncated to 1 nucleotide as described above. Reads (total or separate forward and reverse read counts) were quantitated in running windows as specified in the relevant figure legends before export for plotting using R v4.0.0 in RStudio using the *tidyverse* package [81, 82]. For displaying read counts, values were plotted at the centre of the quantification window and displayed as a continuous line. For RFD plots, RFD values were calculated and plotted as either dots (individual samples) or as a continuous line (multiple sample display) for each quantification window using the formula RFD = (R-F)/(R + F), where F and R relate to the total forward and reverse read counts respectively. The R code to generate these plots can also be found here: https://github.com/FelixKrueger/TrAEL-seq.

A note on read polarity: As a consequence of experimental design, the Illumina sequencing read is the reverse complement of the 3’ extended DNA to which TrAEL adaptor 1 was ligated, and so the first nucleotide of the read is the reverse complement of the last nucleotide 5’ of the break site. To minimise potentially confusing strand inversions, we did not invert the reads during the analysis. In contrast, Sriramachandran *et al*. reversed the polarity of all reads in the analysis pipeline for GLOE-seq [36], which explains the differences in polarity between equivalent analyses in that study and this study.

All sequencing data is available from GEO accession GSE154811.

## Acknowledgements

We thank Paula Koko Gonzales and Nicole Forrester of the Babraham Institute Next Generation Sequencing facility for data generation, Scott Keeney for sharing unpublished data, Adele Marston and Aziz El Hage for yeast strains, and New England Biolabs technical support for helpful answers to a wide range of enzymology questions during the development of this method.

---

**Figure S1:**
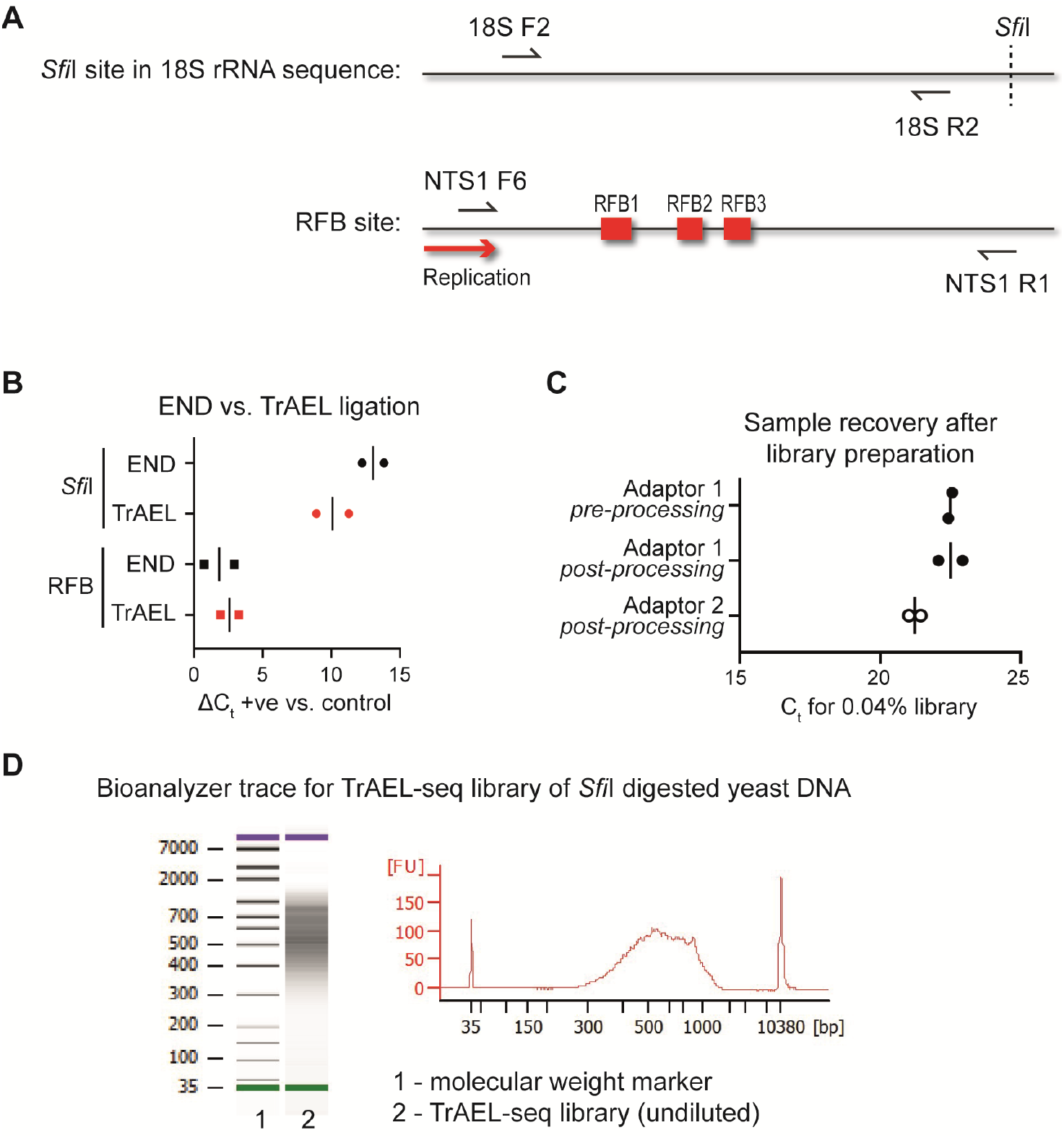
TrAEL-seq library construction details. **A**: Locations of primers used to analyse ligation products at the 18S *Sfi*I site and at the rDNA RFB. Primer sequences are given in Table S2. **B**: Comparison of ligation efficiency for TrAEL and END methods. Adaptors were added according to the TrAEL-seq and END-seq protocols, but DNA was not further processed. qPCR reactions were then performed using the Illumina Read Primer (IRP, which binds to the adaptor in TrAEL and END methods) and 18S or RFB specific primers. For *Sfi*I, ΔC_t_ was calculated comparing reactions with IRP and 18S F2 to reactions with IRP and 18S R2. For RFB, ΔC_t_ was calculated comparing reactions with IRP and NTS1 F6 to reactions with IRP and NTS1 R1 long. **C**: Material recovery after TrAEL-seq processing. For two TrAEL-seq preparations from *Sfi*I-digested genomic DNA that were not performed in parallel, samples were firstly taken directly after gel extraction following TrAEL-seq Adaptor 1 ligation and secondly after pooling the two USER eluates just prior to final amplification. Adaptor 1 qPCR reactions were performed using IRP and 18S F2 (which detects successful ligations to the cleaved *Sfi*I site), Adaptor 2 qPCR reactions used 18S R2 and NEBNext test oligo R and were only performed on the material harvest after USER treatment. The Adaptor 2 PCR product spans the length of the TrAEL fragment from near the *Sfi*I site, across the undefined junction site resulting from sonication and into TrAEL Adaptor 2. **D**: Bioanalyzer trace for the amplified library of *Sfi*I-digested yeast genomic DNA. 1 μl of the 10.5 μl final library was run on a DNA high sensitivity Bioanalyzer chip. This shows a complete absence of adaptor or primer dimers, which is only achieved after two successive AMPure purifications. This trace is typical for TrAEL-seq libraries.

**Figure S2:**
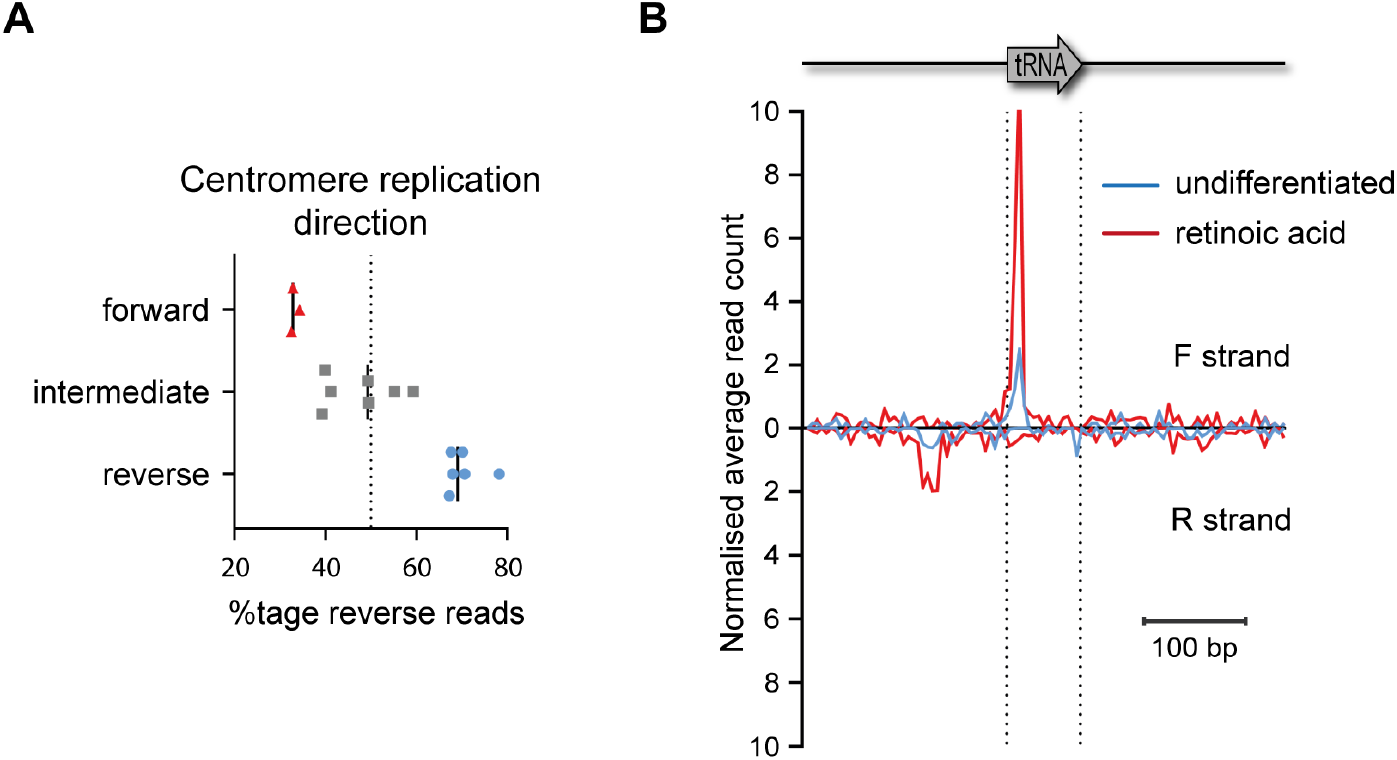
Additional data for detection of replication fork stalling by TrAEL-seq. **A:** Replication direction of centromeres, calculated based on the *cdc9*-AID GLOE-seq data (SRA accession: SRX6436838). Percentage of reverse reads was determined in the regions −1000 to −500 bp and +500 to +1000 bp relative to the annotated centromere, and the average of these values plotted. The region from −500 to +500 bp was excluded as replication fork stalling in this region obscures the replication direction. **B:** Average TrAEL-seq profiles across tRNAs +/- 200 bp for undifferentiated and retinoic acid-treated hESC cells. Reads are separated by orientation on forward or reverse strands, all tRNAs are included.

**Figure S3:**
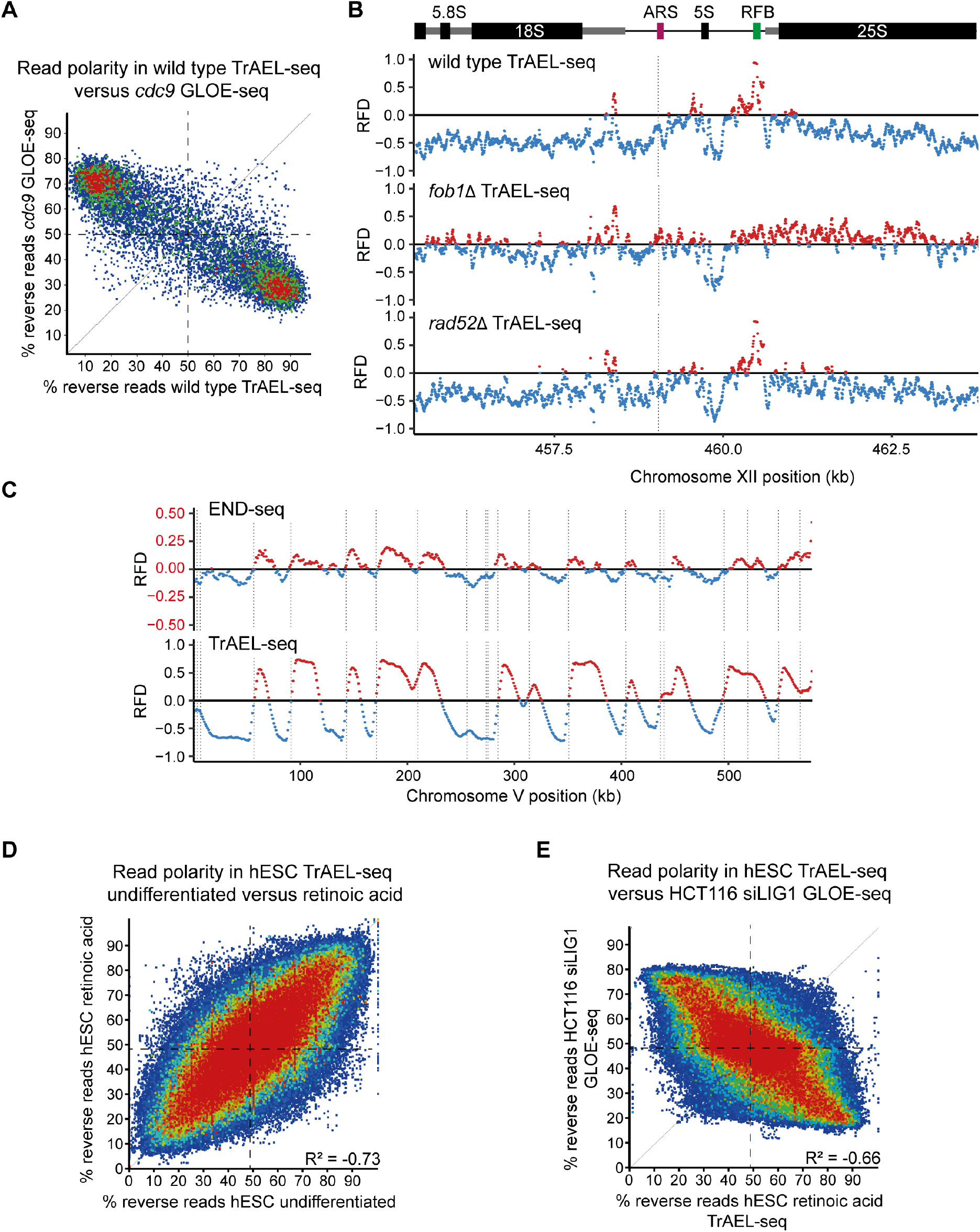
Additional data for replication fork directionality of TrAEL-seq data. **A**: Scatter plot showing the percentage of reverse reads compared to all reads in 1 kb genomic windows spaced every 1 kb, comparing TrAEL-seq data from wild-type cells and GLOE-seq data from Cdc9-depleted cells (SRA accession: SRX6436838). **B**: RFD plots showing TrAEL-seq data for wild type, *fob1*Δ and *rad52*Δ across a single rDNA repeat. The 35S rRNA gene transcribed by RNA polymerase I is shown as a thicker grey line and is transcribed right-to-left in this representation. Mature rRNA genes are shown in black, the RFB and the ARS are also annotated. **C**: RFD plot comparing END-seq and TrAEL-seq data generated from two halves of an agarose plug containing 10 million wild type *3xCUP1* cells grown in synthetic complete glucose media. Note that the scale for the END-seq data is expanded as the bias in read polarity is much smaller in END-seq libraries. **D**: Scatter plot showing the percentage of reverse reads compared to all reads in 250 kb genomic windows spaced every 10 kb, comparing TrAEL-seq data from undifferentiated and retinoic acid treated hESC cells. **E:** Scatter plot showing the percentage of reverse reads compared to all reads in 250 kb genomic windows spaced every 10 kb, comparing TrAEL-seq data from retinoic acid treated hESC cells to GLOE-seq data from LIG1-depleted HCT116 cells (average of SRA accessions: SRX7704535 and SRX7704534).

**Figure S4:**
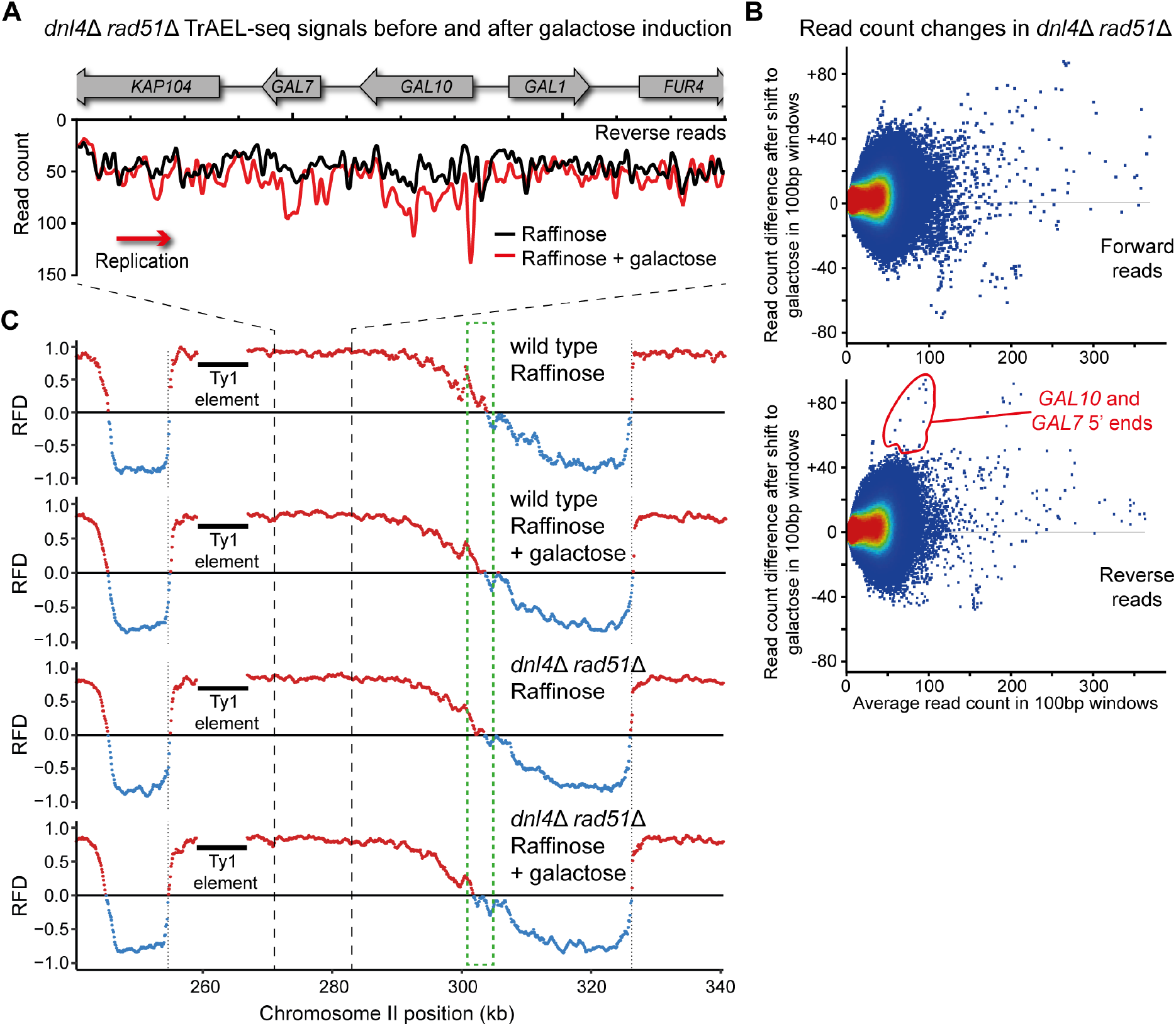
Additional data for detection of environment-dependent replication differences. **A**: Plot of read count differences across GAL locus on galactose induction for *dnl4*Δ *rad51*Δ mutant, as Fig. 4B. **B**: MA plots of changing read count across the genome on galactose induction for *dnl4*Δ *rad51*Δ mutant, as Fig. 4C. **C**: RFD plots showing replication profile of the region surrounding the GAL locus with and without galactose induction. Green box shows the site at which the replication fork which passes through the GAL locus encounters the oncoming fork from ARS211.

**Figure S5:**
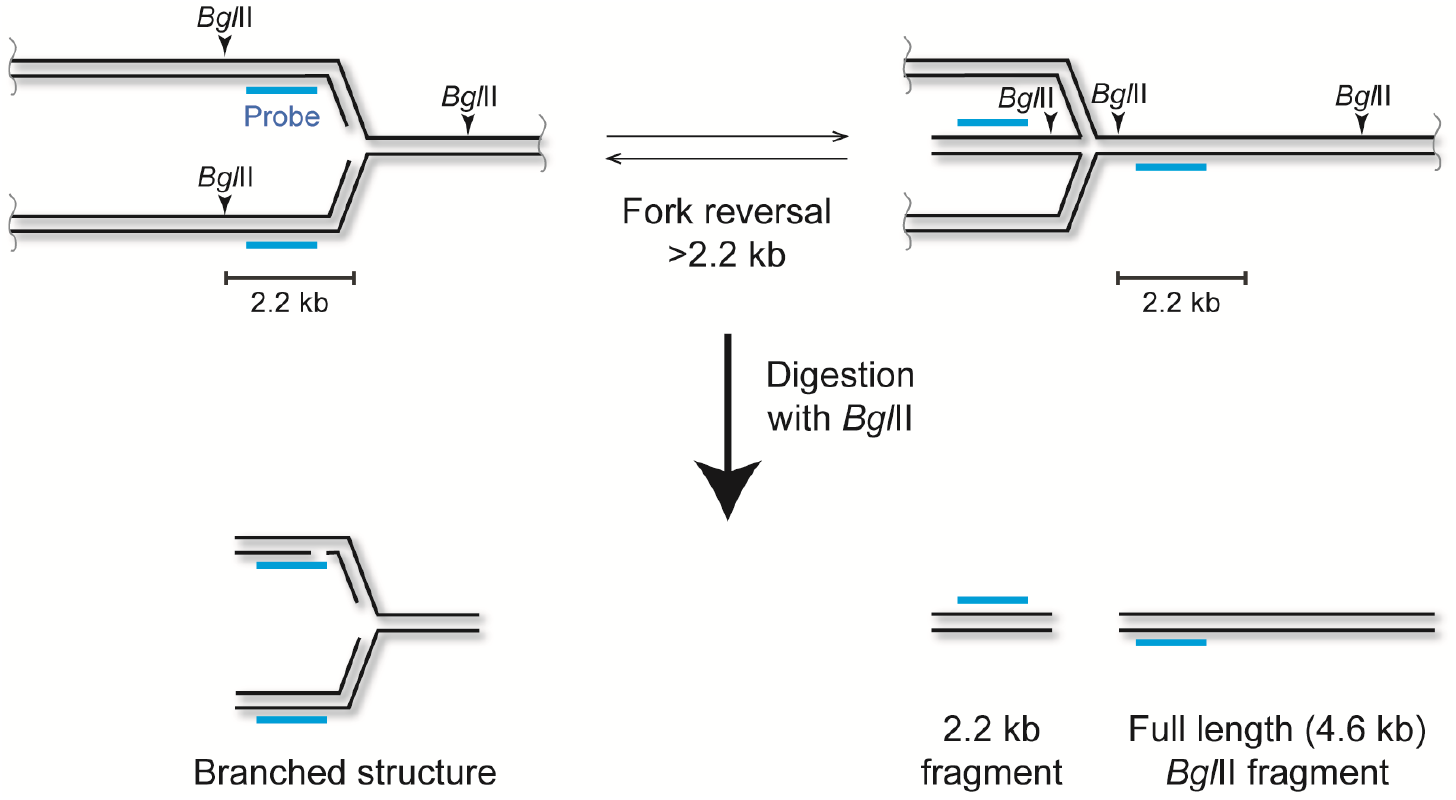
Means by which reversed forks could resemble DSBs in southern analysis. All southern blot analyses that have reported direct detection of DSBs at RFBs utilise a restriction digestion to separate the region of interest. For the yeast RFB, to our knowledge the enzyme used has always been *Bgl*II, the cleavage sites for which lie 2.2 kb and 2.4 kb each side of the RFB. Forks that reverse past the *Bgl*II site would yield a *Bgl*II fragment the same size (2.2 kb) as a fork that is cleaved at the RFB. Only fragments that would hybridise to the probe (blue) are shown.

## Tables

**Table S1:** Table of yeast strains used in this study.

| Name | Genotype | Reference |
| --- | --- | --- |
| BY4741 | MATa *his3Δ1 leu2Δ0 met15Δ0 ura3Δ0* | [84] |
| *fob1Δ* | MATa *his3Δ1 leu2Δ0 met15Δ0 ura3Δ0 fob1::HygMX6* | Gift from Aziz EL Hage |
| *rad52Δ* | MATa *his3Δ1 leu2Δ0 met15Δ0 ura3Δ0 rad52::KanMX4* | [85] |
| *rnh201Δ* | MATa *his3Δ1 leu2Δ0 met15Δ0 ura3Δ0 rnh201::KanMX4* | [85] |
| *rnh202Δ* | MATa *his3Δ1 leu2Δ0 met15Δ0 ura3Δ0 rnh202::KanMX4* | [85] |
| *clb5Δ* | MATa *his3Δ1 leu2Δ0 met15Δ0 ura3Δ0 clb5::KanMX4* | [85] |
| *dnl4Δ rad51Δ* | MATa *his3Δ1 leu2Δ0 met15Δ0 ura3Δ0 dnl4::KanMX4 rad51::NatMX6* | This study |
| *3xCUP1* | MATα *his3Δ1 leu2Δ0 lys2Δ0 ura3Δ0 ade2Δ::LEU2 cup1Δ::ADE2-3xCUP1* | [72] |
| SK1 *dmc1Δ* diploid | MATa/α *ho::LYS2 lys2 ura3 leu2::hisG his3::hisG trp1::hisG dmc1::NatMX6/dmc1::KanMX6* | This study |

**Table S2:**
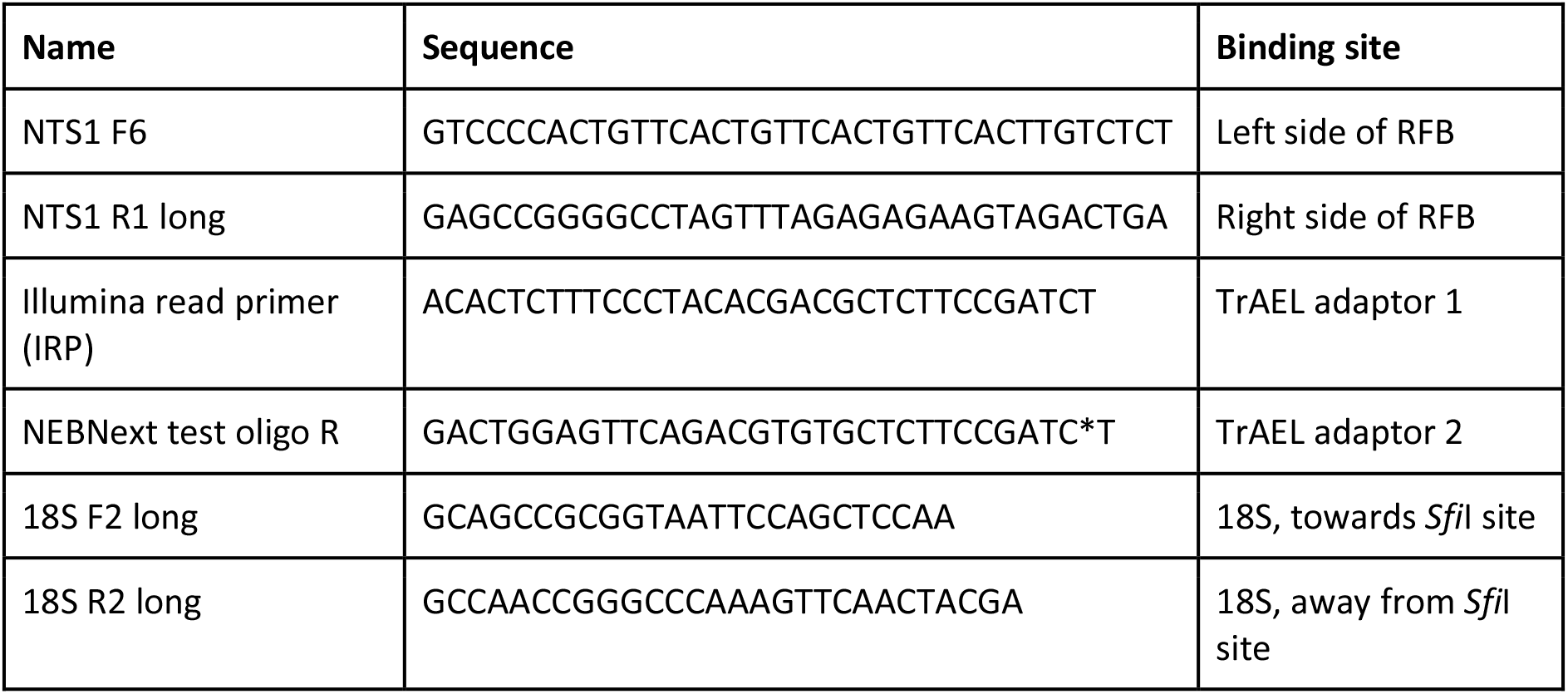
Table of oligonucleotides used in this study.

